# Minos: variant adjudication and joint genotyping of cohorts of bacterial genomes

**DOI:** 10.1101/2021.09.15.460475

**Authors:** M. Hunt, B. Letcher, K.M. Malone, G. Nguyen, M.B. Hall, R.M. Colquhoun, L. Lima, M.C. Schatz, S. Ramakrishnan, CRyPTIC consortium, Z. Iqbal

**Author notes:** Please see later section “CRyPTIC consortium” for details.

## Abstract

Short-read variant calling for bacterial genomics is a mature field, and there are many widely-used software tools. Different underlying approaches (eg pileup, local or global assembly, paired-read use, haplotype use) lend each tool different strengths, especially when considering non-SNP (single nucleotide polymorphism) variation or potentially distant reference genomes. It would therefore be valuable to be able to integrate the results from multiple variant callers, using a robust statistical approach to “adjudicate” at loci where there is disagreement between callers. To this end, we present a tool, Minos, for variant adjudication by mapping reads to a genome graph of variant calls. Minos allows users to combine output from multiple variant callers without loss of precision. Minos also addresses a second problem of joint genotyping SNPs and indels in bacterial cohorts, which can also be framed as an adjudication problem.

We benchmark on 62 samples from 3 species (*Mycobacterium tuberculosis, Staphylococcus aureus, Klebsiella pneumoniae*) and an outbreak of 385 *M. tuberculosis* samples. Finally, we joint genotype a large *M. tuberculosis* cohort (N*≈*15k) for which the rifampicin phenotype is known. We build a map of non-synonymous variants in the RRDR (rifampicin resistance determining region) of the *rpoB* gene and extend current knowledge relating RRDR SNPs to heterogeneity in rifampicin resistance levels. We replicate this finding in a second *M. tuberculosis* cohort (N*≈*13k).

Minos is released under the MIT license, available at https://github.com/iqbal-lab-org/minos.

## Introduction

The use of whole genome short-read sequence data to study cohorts of bacterial genomes from a single species is now standard practise (eg [1, 2]). There are a multitude of variant callers, which analyse reads from a sample and make statements about where it differs from a fixed reference genome. However there is no one best variant caller, or even approach - all have strengths and weaknesses. The single-sample-inference problem is well studied and understood – mapping to a reference works well where the sample and reference are reasonably close, and primarily for SNPs (SAMtools[3], snippy https://github.com/tseemann/snippy), but fares progressively worse as the reference and sample diverge[4]. On the other hand, methods based on local assembly (GATK[5], Octopus[6]) are better able to detect small indels, and those based on global assembly (Cortex[7], McCortex[8]) are exceptionally specific, robust to reference-choice, and better at accessing clustered SNPs or indels up to a few kb in size, but at a cost in sensitivity. Ideally, it would be valuable to be able to combine the output of two different methods (“callsets”) in some rigorous manner, resulting in a product better than either. Simply using the union of callsets gives no control over false discovery rate, and the intersection is too conservative, losing the benefit of callers with different strengths. This “variant adjudication” problem of rigorously combining callsets, where discordances between input variants are resolved and then variant sites are genotyped, is the first challenge we address in this study.

Moving beyond single samples, there are many use-cases where one needs to jointly analyse a cohort, producing a matrix of variants versus samples and making binary or probabilistic statements (genotype calls) at all positions that are segregating in the cohort. This is more tricky, primarily because the density of variation increases with cohort size, and inevitably there are situations where SNPs and indels overlap. This is typically described as joint genotyping[5], and itself is a form of adjudication problem – once we have the full list of segregating sites and alleles, we can revisit all samples and adjudicate each one. This is the second main problem we address in this study.

Finally, underlying both, there is an important technical challenge: how to combine Variant Call Format (VCF)[9] files, either for the same sample from different variant callers, or from multiple samples in a cohort when collecting a list of segregating sites. In both cases, we want a clean VCF file with a set of non-overlapping records, each representing a segregating site with alternate alleles, which can be independently genotyped (see Figure 1). This likely entails combining independent overlapping variants from the input VCF files into some consistent multi-allelic record. This particular problem is technically awkward, and can in theory get arbitrarily ugly - in the pathological worst case there could be overlapping records that gradually tile across the whole genome.

**Figure 1:**
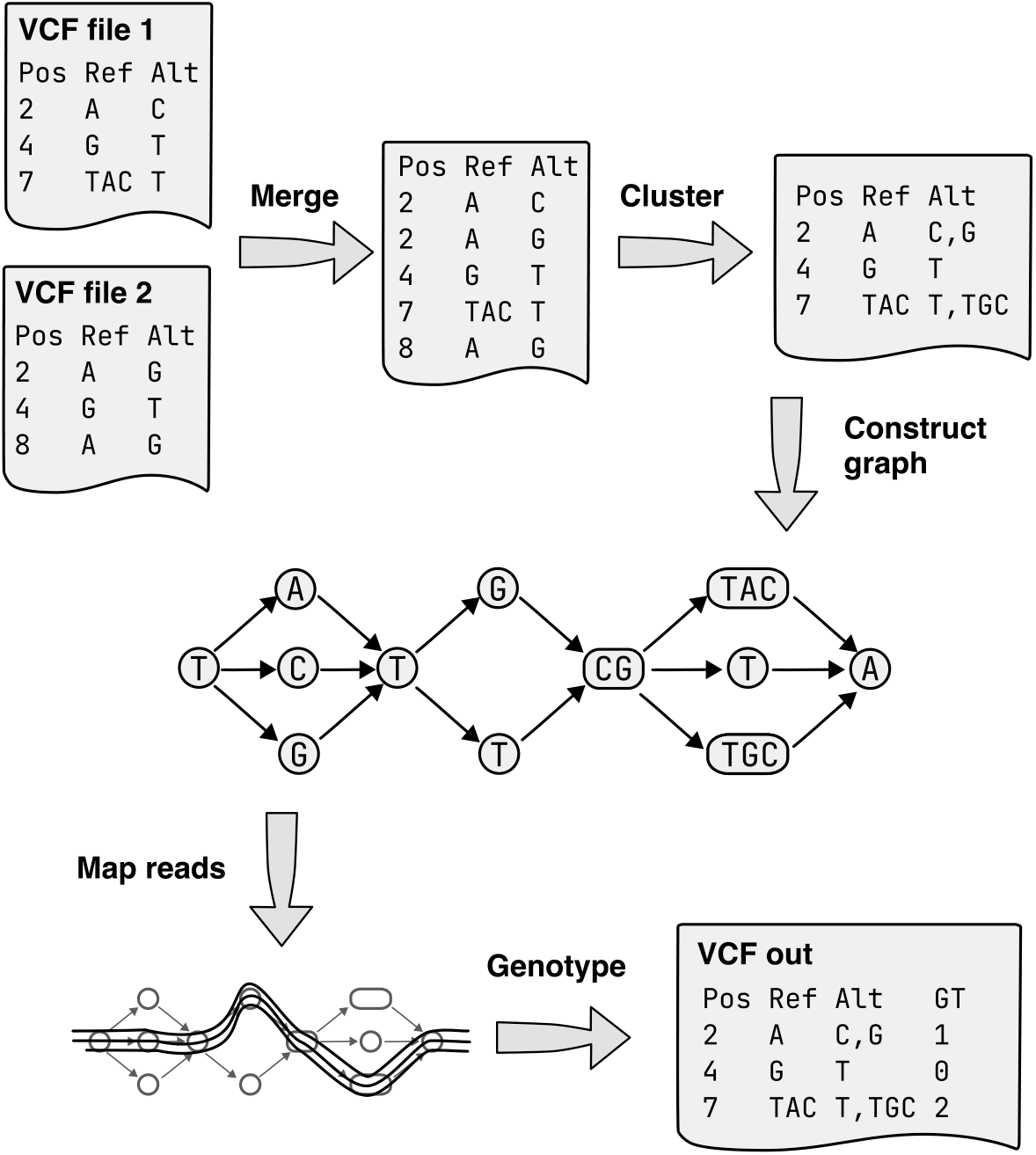
Variant adjudication pipeline implemented by Minos. Input variants in one or more VCF file(s) are merged to make a deduplicated set of variants. When running on a single sample, the input VCF files could be from different tools. When joint genotyping across samples, there is one VCF file originating from each sample. Next, overlapping variants are clustered together – for example the variants at positions 7 and 8 – allowing the construction of a non-nested variation graph. Genotype calls are made using read mapping to the graph.

Our motivation was the desire to study tens of thousands of *Mycobacterium tuberculosis* genomes for the CRyPTIC project[10]. This project sequenced and phenotyped over 15,000 isolates for resistance to 13 different drugs using a 96-well microtitre plate[11], and then applied methods such as Genome-Wide Association Studies (GWAS) to analyse the genetic basis for drug resistance. *M. tuberculosis* has relatively low levels of diversity by bacterial standards, with no recombination, relatively few mobile genetic elements, and a small pan-genome[12]. However, despite this, almost 17% of the 4.4Mb genome was variable within the cohort, and in some regions almost every single base harboured a multiallelic SNP or indel. Typical approaches tuned for high precision callsets would refuse to make calls in such dense regions, but for our purposes we needed to be able to both represent and correctly genotype these regions.

There has been prior work on these problems, with recent tools focussed on human data. Joint genotyping is available in GATK, however it relies on machine-learning-based filtering (VQSR) generated from human-specific truth-data. Applying GATK to non-human species required considerable efforts to train a black box VQSR for each new species (eg see [13] for *Plasmodium*). At the time of writing, GATK explicitly does not support bacterial data, although a new version for this is in development. There are also graph-based genotypers for structural variants that operate on similar principles to ours, although with different graph structures and genotyping models, such as Paragraph[14] and vg[15].

Two other graph-mapping based tools are available: BayesTyper[16] maps reads to a directed acyclic graph of informative kmers, and GraphTyper[17] maps to local graphs of SNPs and indels from pre-mapped reads. Although it is possible to use these tools as an adjudicator, consuming more than one VCF file and outputting a single VCF file, they were developed for human data and do not scale to our use case of tens of thousands of bacterial samples. In particular, neither tool properly combines conflicting and/or overlapping variants from the input callers, so that the user is faced with contradictory output. This is a non-trivial development problem, as described above. Thus, although they use graphs and mapping, neither tool directly seeks to solve our problem.

Our approach was to build a pure adjudicator, able to run a single command that can take multiple VCF files, handle all overlapping variants, and output a single accurate callset with no inconsistencies. Intuitively, read pileup can be used to test goodness-of-fit. Reads should map perfectly to a reference containing the correct allele, so comparing pileups on alternate alleles can resolve disagreements between callsets. Of course, it would be prohibitively expensive to remap all reads to every input allele independently and then compare the pileup on each allele. Instead, we build a genome graph of the combined alleles from all callers and map once to that, using gramtools[18]. Reads naturally align to the correct allele, and we can genotype using the resulting coverage and ambiguity information (described below). We implemented this, plus a workflow for joint genotyping cohorts, in our new tool, Minos.

We first use 62 high quality polished long-read assemblies from three bacterial species and benchmark Minos against BayesTyper and GraphTyper. We then apply all three tools to an *M. tuberculosis* outbreak (*N* = 385) in the UK in 2013, evaluating precision for both reference and non-reference calls.

Finally, we apply our method to our motivating problem – studying antimicrobial resistance in large cohorts of *M. tuberculosis*. We first joint-genotype the CRyPTIC global cohort of *M. tuberculosis* genomes (*N* = 15, 215) which contains around 700,000 variants (roughly one SNP every 5bp). We focus on the 81bp rifampicin-resistance determining region (RRDR) in the *rpoB* gene, which bears an enormous level of variation. The WHO-endorsed Xpert^®^ MTB/RIF assay assumes any non-synonymous SNP in this region causes resistance to rifampicin[19], although as we discuss below, the story is a little more complex. We restrict to non-synonymous SNPs and indels, and give an unprecedented map of dense variation in the RRDR and how strongly each variant correlates with resistance. We find the five known “borderline” mutations[20] but also show that there are more. We then joint-genotype a second, independent, cohort of 13,411 *M. tuberculosis* genomes which have also been phenotyped for rifampicin, and replicate the finding. We consider the significance of these findings in the Discussion.

## Results

We developed a new tool called Minos, which takes putative variant calls as input, adjudicates between all of the calls, and reports a final accurate callset. It uses the standard Variant Call Format (VCF) for its input and output. Additionally, Minos includes a Nextflow[21] pipeline to joint genotype large numbers (tens of thousands) of samples, producing a set of calls at the same variant sites across all samples. See Figure 1 for an overview of the pipeline, and the Methods section for a complete description.

To evaluate the performance of Minos compared to other tools, we also developed a new tool called Varifier, which calculates the precision and recall of a variant callset in a novel way that accounts for insertions and deletions in addition to SNPs. As described in full in the Methods section, it compares each allele against a “truth” genome, assumed to contain no errors, to determine if each genotype call is a true or false positive. For each sample, we used a truth genome assmbled from long reads (PacBio or Oxford Nanopore), and polished using the same Illumina reads that were used for variant calling. We used Varifier and these truth genomes to benchmark Minos against GraphTyper and BayesTyper with simulated and real data.

### Single-sample benchmarking

We performed an initial sanity check that BayesTyper, GraphTyper, and Minos all work as expected, using simulated reads made with ART[22] and simulated variants in the *M. tuberculosis* H37Rv reference genome[23]. All tools performed near perfectly on this simple data set, affirming that they make no major errors (Supplementary Figure 1, Supplementary Tables 1,2). The default Minos filters dropped the recall of SNPs and short indels slightly (Supplementary Figure 1), due to unrealistically low variation in read depth in the simulations, which caused the “MIN GCP” filter (see Methods) to fail true positive calls. However, as shown later, this is not an issue in real sequencing read data. Having confirmed all tools passed a basic test, we move on from the simulations to empirical data.

Next, the tools were compared using real data from *M. tuberculosis, Staphylococcus aureus*, and *Klebsiella pneumoniae*, using samples which each had a high quality polished long read assembly to act as truth, and matched Illumina data (see Methods). We selected reference genomes for each species (1 for *M. tuberculosis*, 2 for *S. aureus* and 5 for *K. pneumoniae*) to reflect the diversity of the species.

For each Illumina data set, reads were trimmed using Trimmomatic[24], mapped to the reference genome with BWA MEM[25], and PCR duplicate reads were removed with SAMtools. Variants were called independently using two variant callers with orthogonal strengths: SAMtools/BCFtools is pileup-based, with high sensitivity for SNPs and low precision for indels; Cortex is assembly-based, with high precision for SNPs and indels, but lower recall. The SAMtools/BCFtools and Cortex callsets were input to BayesTyper, GraphTyper and Minos, resulting in a single set of calls from each tool (BayesTyper and GraphTyper required additional processing of the SAMtools/Cortex VCF files, described in the supplementary Methods). In order to maximise recall, and because the adjudication tools should remove false-positive variants, the unfiltered callsets from SAMtools and Cortex were used. All results shown are using the default variant call filters for each tool, except where noted, and with unreliable regions of the genomes masked.

Note that GraphTyper can be run in two genotyping modes: default and “sv” (described in [26]). The “sv” mode resulted in significantly worse results in many cases (Supplementary Table 3), therefore it is not discussed further in this manuscript. All GraphTyper results in this manuscript refer to running in default mode.

The results are summarised in Table 1 (also Supplementary Figures 2, 3). Minos achieved the best F-score and recall across seven of the eight data sets. The biggest variation between tools was seen in the recall. Although GraphTyper had the highest precision, Minos was equally precise in three of the data sets, and the biggest difference in mean precision between Minos and GraphTyper was 0.05%.

**Table 1:** Mean precision, recall and F-score on each empirical data set with each reference genome. Numbers in bold show the best precision, recall and F-score for each species and reference genome.

| Species | Number of samples | Reference genome | Tool | Mean precision | Mean recall | Mean F-score |
| --- | --- | --- | --- | --- | --- | --- |
| *M. tuberculosis* | 17 | H37Rv | BayesTyper | 0.9995 | **0.9217** | **0.9579** |
|  |  |  | GraphTyper | **0.9997** | 0.8938 | 0.9422 |
|  |  |  | Minos | **0.9997** | 0.9181 | 0.9559 |
| *S. aureus* | 28 | TW20 | BayesTyper | 0.9984 | 0.8669 | 0.9279 |
|  |  |  | GraphTyper | **0.9990** | 0.7530 | 0.8545 |
|  |  |  | Minos | 0.9988 | **0.8786** | **0.9347** |
|  |  | USA300 | BayesTyper | 0.9993 | 0.8671 | 0.9283 |
|  |  |  | GraphTyper | **0.9995** | 0.7506 | 0.8534 |
|  |  |  | Minos | 0.9994 | **0.8792** | **0.9353** |
| *K. pneumoniae* | 17 | GCF_000784945.1 | BayesTyper | 0.9990 | 0.9052 | 0.9495 |
|  |  |  | GraphTyper | **0.9999** | 0.9063 | 0.9505 |
|  |  |  | Minos | **0.9999** | **0.9143** | **0.9550** |
|  |  | GCF_001952915.1 | BayesTyper | 0.9995 | 0.8800 | 0.9346 |
|  |  |  | GraphTyper | **0.9999** | 0.8788 | 0.9340 |
|  |  |  | Minos | 0.9994 | **0.8922** | **0.9417** |
|  |  | GCF_003073315.1 | BayesTyper | 0.9996 | 0.9267 | 0.9617 |
|  |  |  | GraphTyper | **0.9999** | 0.9297 | 0.9634 |
|  |  |  | Minos | 0.9998 | **0.9367** | **0.9672** |
|  |  | GCF_003076555.1 | BayesTyper | 0.9994 | 0.9397 | 0.9686 |
|  |  |  | GraphTyper | **0.9999** | 0.9387 | 0.9683 |
|  |  |  | Minos | **0.9999** | **0.9438** | **0.9710** |
|  |  | GCF_011006575.1 | BayesTyper | 0.9995 | 0.9078 | 0.9511 |
|  |  |  | GraphTyper | **0.9999** | 0.9075 | 0.9513 |
|  |  |  | Minos | 0.9998 | **0.9238** | **0.9602** |

### Performance

On the real bacterial data, all tools had a relatively fast run time and low RAM usage (Supplementary Table 4, Supplementary Figure 4). On each data set, GraphTyper had the shortest run time and smallest RAM usage, followed by Minos and then BayesTyper. The median wall clock time for Minos across the data sets ranged from approximately 2.5 minutes per sample (*M. tuberculosis*) to 6 minutes (*K. pneumoniae*). *K. pneumoniae* required the highest RAM, where the median peak usage across all runs for Minos was 2.5GB.

### Joint genotyping of cohorts

We wrote a Nextflow pipeline, included in the Minos code repository, to easily joint genotype large numbers of samples. It outputs a single VCF file containing all samples genotyped at the same sites, and the same information in per-sample VCF files. We tested this pipeline, together with BayesTyper and GraphTyper, to re-analyse 385 *M. tuberculosis* [27] samples from an outbreak in the UK, which we call the “Walker 2013” data set. Note that BayesTyper and GraphTyper are not set up for this use case, and therefore processing the data was required to attempt to use these tools in this manner, as described in the Methods section.

Our final analysis was to joint genotype two large *M. tuberculosis* cohorts of more than ten thousand samples each, using the results to analyse the phenotypes and corresponding genotypes in the 81bp rifampicin resistance determining region (RRDR) of the *rpoB* gene. The first large data set (“CRyPTIC”) consisted of 15,215 samples released by the CRyPTIC project[10], phenotyped using a microtitre plate assay[28, 11], and the second (“Mykrobe”) data set consisted of 13,411 samples previously published[29] and phenotyped using traditional culture-based DST (drug susceptibility testing).

Variants were called on each sample independently with the same methods as above using Cortex and SAMtools, and then calls adjudicated with Minos. Each of these per-sample Minos VCF files was used as input to each of the three tools. The result was a callset for each tool, at the same variant sites for all samples for that tool (variant sites were not the same between tools). Minos is the only tool of the three that is set up to consistently merge all input VCF records, so that no two sites in the output contain reference positions in common. Furthermore, this means Minos does not, unlike BayesTyper and GraphTyper, output two separate VCF records with incompatible genotype calls. This is discussed in more detail in the Supplementary Material.

A summary of the data sets and variants output by Minos is given in Table 2. A typical *M. tuberculosis* genome might be around 1000 to 2000 SNPs distant from the H37Rv reference genome. Since these are spread across a 4.4Mb genome, excessive density of variants is not a problem for the single-sample adjudication problem. However when joint genotyping the larger cohorts, we are adjudicating all segregating variation, covering 17-18% of the genome, including some regions of hundreds of base-pairs where almost every single base is a multiallelic SNP or indel. These dense regions present a scaling challenge to graph genome algorithms, depending on implementation and indexing strategy.

**Table 2:**
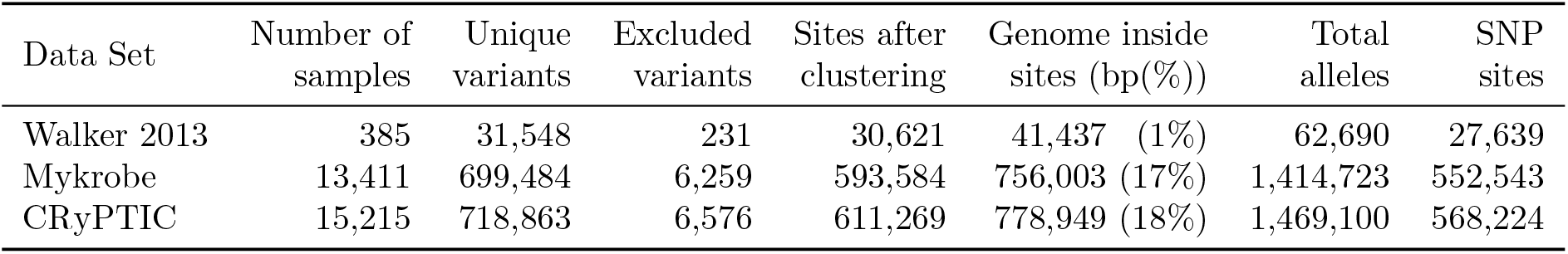
Summary of *M. tuberculosis* data sets used for joint genotyping. “Genome inside sites” is the total length of all reference alleles across all sites after clustering. It is reported as the total number of base pairs, and in parentheses as a percentage of the 4.4Mbp H37Rv reference genome. SNP sites is the number of sites where all alleles have length 1.

| Data Set | Number of samples | Unique variants | Excluded variants | Sites after clustering | Genome inside sites (bp(%)) | Total alleles | SNP sites |
| --- | --- | --- | --- | --- | --- | --- | --- |
| Walker 2013 | 385 | 31,548 | 231 | 30,621 | 41,437 (1%) | 62,690 | 27,639 |
| Mykrobe | 13,411 | 699,484 | 6,259 | 593,584 | 756,003 (17%) | 1,414,723 | 552,543 |
| CRyPTIC | 15,215 | 718,863 | 6,576 | 611,269 | 778,949 (18%) | 1,469,100 | 568,224 |

### Outbreak analysis

We evaluated performance of Minos and alternate tools on genomes in the Walker 2013 data set, from an outbreak of *M. tuberculosis*. To measure the precision and recall of the three tools before and after joint genotyping, the 17 *M. tuberculosis* samples used earlier were added into the outbreak data set. Joint genotyping the samples generally had a negligible effect on precision and recall (Supplementary Table 5). BayesTyper recall increased by 0.07%, whereas the recall of GraphTyper and Minos dropped by 4.2% and 0.6% respectively. BayesTyper precision fell by 0.13%, and both GraphTyper and Minos increased by 0.01%.

Up to this point in the manuscript, precision has always been calculated by considering only non-reference allele calls, focussing on the differences between a given sample and the reference genome. However, joint genotyping involves genotyping every sample at every variant site. As a result, the majority of calls have the reference genotype, and correctly genotyping these cases is critical for applications such as building a phylogenetic tree, computing a genetic distance matrix or for GWAS. The difference between excluding or including reference calls for each tool is shown in Figure 2 and Supplementary Table 5. After joint genotyping, and including reference genotype calls in the calculation, Minos achieved a precision of 99.95%, compared with 92.87% and 88.85% for BayesTyper and GraphTyper respectively.

**Figure 2:**
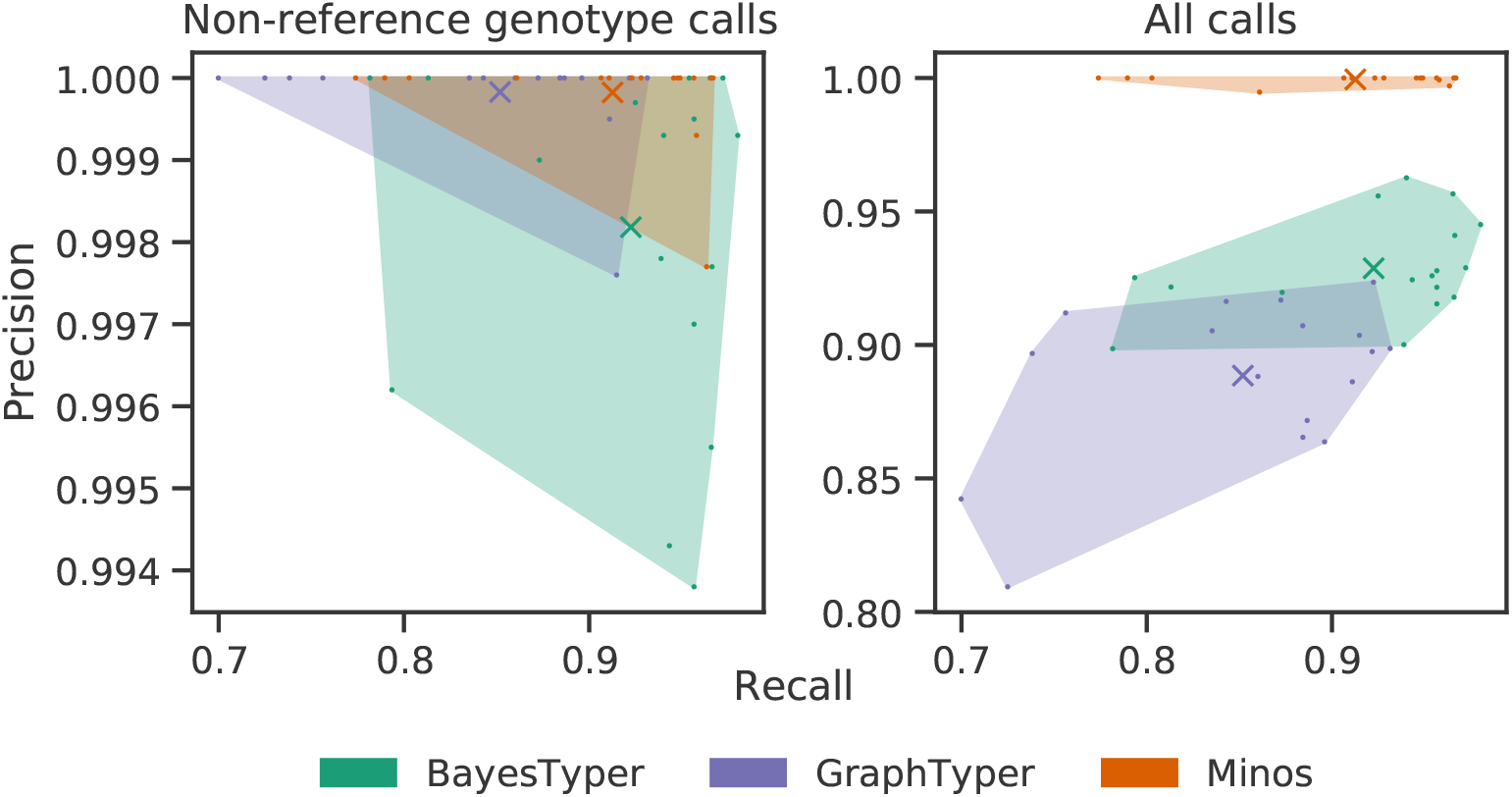
Precision and recall when joint genotyping *M. tuberculosis* outbreak data. The left plot considers non-reference allele calls only, ie the variant sites that are genotyped to be different from the reference genome. The right plot shows the results when all allele calls are included. Individual samples are marked as dots, and the mean precision and recall for each tool is shown as a cross. The convex hull of the data points for each caller is shaded with an associated colour.

### Association of rifampicin resistance with the RRDR region of *rpoB*

Returning to the motivating problem for which Minos was developed, we applied Minos to two large *M. tuberculosis* cohorts (“CRyPTIC” and “Mykrobe” data sets) for which we have associated resistance phenotype data for the first-line drug rifampicin. We sought first to confirm that Minos would indeed function at this scale, and then to build a detailed map of variation in the RRDR (rifampicin resistance determining region) of the *rpoB* gene. Minos output genotype calls at 593,584 and 611,269 variant sites for the Mykrobe and CRyPTIC data sets respectively (Table 2). These sites cover approximately 17-18% of the 4.4Mb H37Rv reference genome.

The total turnaround time of the pipeline, which was limited to a maximum of 2000 concurrent tasks, was approximately 23 hours for the Mykrobe data set and 26 hours for CRyPTIC. These represent real-world times, since the compute cluster we used was shared with numerous other users. A breakdown of the run times and maximum RAM usage is provided in Supplementary Table 6. The pipeline is optimised to minimise memory, with the peak memory used when merging and clustering variants at less than 8GB. The majority of the run time comprises running Minos on each sample, which required less than 2GB of RAM per sample.

We then used the joint genotyping output to analyse the genotype-phenotype relationship within the RRDR of the *rpoB* gene, by restricting to the 13,259 samples of the Mykrobe data set, and the 8,955 samples of the CRyPTIC data set with a high quality rifampicin phenotype[10] (from 12,099 CRyPTIC samples with any quality rifampicin phenotype). Figure 3 shows all the identified amino acid variants – substitutions, insertions and deletions – plotted along the RRDR, with their prevalence and proportion of resistant samples for the CRyPTIC data. The same plots for the Mykrobe data and all CRyPTIC data are given in Supplementary Figure 5, and the raw data are in Supplementary Table 7.

**Figure 3:**
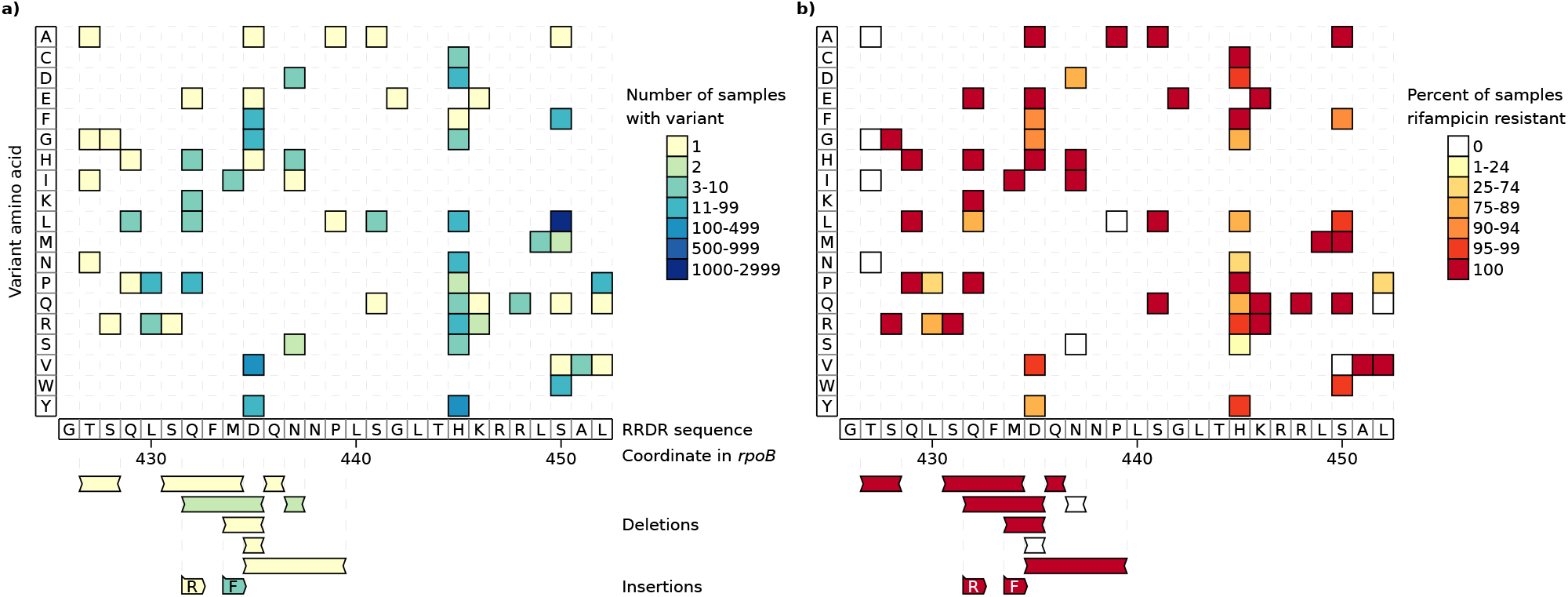
All amino acid variants identified in the RRDR of the *rpoB* gene by joint genotyping 8,955 samples from the CRyPTIC *M. tuberculosis* data set. Each plot shows the RRDR region from left to right. Single amino acid variants are shown in the upper grid, with the y axis corresponding to the variant amino acid. The lower area shows deletions and insertions, with the inserted sequence given in the coloured boxes. For example, the leftmost deletion of amino acids TS at position 427-428 is found in one sample, which is resistant. The leftmost insertion adds R after the S at position 431 (found in one resistant sample). The plots show the same variants, but with different colour schemes. In a) each variant is coloured by the number of samples possessing that variant. Plot b) colours the variants by the percent of samples with that variant that are rifampicin resistant.

As expected the variant S450L, found in 1,919 of the samples, dominates. The next most common variant D435V appears in 345 samples. Several rare indels were identified, most of which appear to cause rifampicin resistance. In the CRyPTIC samples, two insertions were identified (R inserted at 431-432 and F at 433-434), and all nine samples with either of these are rifampicin resistant. Six of the eight deletions arise only in resistant samples. In the Mykrobe data there were seven deletions and three insertions, all of which were only found in resistant samples.

We find a total of 72 distinct amino acid mutations and indels within the 81bp RRDR in the CRyPTIC samples with high quality phenotypes, including the 5 mutations classified as “borderline resistant” by the WHO[20] (H445L, H445N, D435Y, L452P, L430P). There are 13 variants in the CRyPTIC samples where the number of susceptible samples is greater than the number of resistant samples. The most common are amongst the borderlines listed above – L430P, where 45/67 samples are susceptible (and 9/19 susceptible in the Mykrobe data), and H445N, where 16/22 samples are susceptible (2/6 in Mykrobe samples). The remaining such variants are rare, each seen in up to five samples. Full counts can be seen in Supplementary Figure 5. Even if the known borderline mutations are excluded, Figure 3 (right panel) shows a large number of moderate to low frequency variants with a range of correlations with resistance. We discuss these results further below.

## Discussion

Variant analysis from short read sequence data is by now a mature field, and there are many tens of different tools for detecting SNPs and short indels[30]. Setting aside performance issues and focussing entirely on completeness and correctness of the inferred SNPs and indels, it is clear that there is no single best tool. The relative weight given to mapping, assembly, pairedend information and species-specific optimisation (eg via machine learning) result in different strengths and weaknesses. This leads to two rational choices: first, to benchmark for your chosen application and choose the best tool, and second to find a way to combine the strengths of different callers. When setting up the CRyPTIC project, we observed that there was no off-the-shelf solution to this problem, and set out to produce an easy-to-use tool that would do this in a rigorous manner. Minos was the result, which we incorporated into a workflow for analysis of *M. tuberculosis* genomes called Clockwork. We found that in terms of single genome analysis, performance was relatively similar to other benchmarked tools – Minos generally had the best recall, and GraphTyper had marginally the best precision. However only Minos would out of the box ingest two (or more) VCF files and output results; the other tools forced users to write code to prepare input data and glue together their processing stages. Combining VCF files with SNPs and indels is particularly challenging for cohorts, where a large proportion of the genome can be variable (17% in our CRyPTIC cohort for example). When analysing the Walker 2013 *M. tuberculosis* outbreak and including reference/wild-type calls, Minos had much higher precision (7-10% higher) than the other tools, and on scaling up, only Minos could process the 12k and 13k CRyPTIC and Mykrobe cohorts.

Rifampicin is a bactericidal drug which is a critical component of the antitubercular arsenal, resistance to which is typically used as an epidemiological proxy for multi-drug resistance (defined as having resistance to both rifampicin and isoniazid), particularly in PCR based rapid diagnostics such as the Xpert^®^ assay. The latest WHO technical report [20] showed that there were 6 known borderline mutations in *rpoB* (of which 5 were in the RRDR)[20, 31, 32, 33], and carefully reported the MIC (minimal inhibitory concentration) distributions for isolates with these mutations. In essence, the distribution of MICs (“the level of resistance”) overlapped with the distribution for wild-type (susceptible) *M. tuberculosis*, which leads to poor reproducibility of binary classification when the threshold between resistant and susceptible (ECOFF) lies in that overlap. Nevertheless, in the light of various reports of worse patient outcome associated with some of these mutations[34, 35,36, 37, 38, 39], the WHO expert group decided that all non-synonymous mutations, even previously unseen ones, should be treated as causing resistance for the purposes of diagnostics and therapy. Our analysis of this, the largest consistently phenotyped cohort to date[10], reveals that the level of resistance caused by mutations in the RRDR is indeed heterogeneous, and reveals further candidate borderline mutations and indels, including a cascade of overlapping rare indels at position 431. This general picture is replicated in the second cohort (Mykrobe data set). For a more nuanced analysis, it is necessary to look at the MIC distribution associated with specific mutations (rather than using a binary resistant/susceptible classification), which has been done in [40].

The main limitation to this study is that for joint genotyping we set a deletion length limit of 50bp. This reflects an underlying design decision: Minos is a tool for combining VCF files from different callers or samples, adjudicating, and outputting an improved VCF file. However VCF is really not an appropriate file format for handling many overlapping small variants and large indels – for example a 5kb deletion covering 20 SNPs. We address this question of how best to genotype and encode multiscale variation (such as SNPs on top of long alternate haplotypes or SNPs under deletions) in a separate study[18]. Nevertheless, we have shown Minos enables users to combine results from their preferred variant callers, integrating their strengths, to reach closer to the underlying truth. It also provides a method for joint genotyping SNPs and indels in bacterial genomes. As genomic analysis is now ubiquitous in bacteriology, we believe Minos will be of wide utility.

## Methods

### Minos pipeline

First we describe the methods used by Minos, which is implemented in Python and available under the MIT license at https://github.com/iqbal-lab-org/minos. The pipeline is outlined in Figure 1. The first two stages process the input variant calls, which must be in one or more VCF files, to produce a single set of calls that can be used to generate a reference graph for read mapping. Initially, calls are normalized and deduplicated to make a single “merged” set of calls. These calls are then “clustered” into variant sites that define the variant graph used for read mapping. The merging and clustering are described below.

#### VCF merging

Each VCF file is processed individually as follows. Variant alleles to be retained for further processing are extracted, where for each record if the genotype (`GT`) field is present then only called the alleles are kept, otherwise all alleles are used. The remaining records and their alleles are written to a new VCF file. Variants are decomposed into unique SNPs and indels using the commands `vcfbreakmulti` and `vcfallelicprimitives -l 10000` from vcflib (https://github.com/vcflib/vcflib), followed by the `normalize` function from vt[41], and finally the `vcfuniq` command from vcflib. These VCF files are loaded into a single data structure containing all the unique variants, plus the origin (ie which VCF file) of each variant.

#### Variant clustering

The merged variants are used to produce a single VCF file of “clustered” variants that is compatible with gramtools, which in turn is used to generate a variant graph and map reads to that graph. Although gramtools supports more complex situations (SNPs on alternate haplotypes separate from the reference genome, SNPs underneath long deletions)[18], we restrict to “non-nested” variation to maintain compatibility with VCF. Therefore overlapping variants must be converted into a single variant site (ie line in a VCF file) containing multiple alleles (an example is given in Supplementary Figure 6). This is straightforward when processing a single sample with a few input VCF files, such as the SAMtools and Cortex VCF files used in this study when benchmarking Minos against other tools. Where possible, all combinations of alleles are generated and included in the graph. However, this is not always feasible when genotyping a large number of samples (hundreds or thousands) because the number of theoretically possible alleles at one site could be very large.

To process a large number of samples, heuristics are used to simplify the variant graph. First, the number of alleles is limited by only allowing deletions of length (by default) *≤*50bp. This prevents a combinatorial explosion where SNPs underneath the deletion can cause impractically large numbers of alleles: *n* biallelic SNPs generates 2^*n*^ alleles. Second, the number of alleles in a variant site is limited to (by default) 5000. If generating all allele combinations at a site results in too many alleles, then only combinations of alleles actually seen in each sample are used. Third, it is possible for the graph to contain the same sequence more than once across multiple variant sites, by choosing different paths through the graph (an example is given in Supplementary Figure 7 - roughly this can happen in low complexity sequence where two alternate deletions of some repetitive sequence can lead to the same final sequence). As each new variant site is added, the previous seven sites are checked and any sites generating duplicate sequences are merged into a single deduplicated site. Removing these duplications is necessary to prevent downstream read mapping issues caused by ambiguous mapping to different paths in the graph that are really the same sequence. Finally, to reduce RAM usage and run time, there is an option to split the graph into chunks, with read mapping run separately on each of these chunks. Using this option requires the reads to be in a sorted indexed BAM file, so that the reads for each chunk can be efficiently extracted for mapping.

#### Graph mapping and genotyping

The clustered VCF file made in the previous stage is input to the `build` command of gramtools[18] to make a variant graph for read mapping. Reads are mapped to the graph using the gramtools command `quasimap`. Each variant site is genotyped using the output from gramtools, which reports the number of reads mapped to each allele, and the read depth across each position of each allele. Minos supports haploid genotyping calling only, using the model described below. Minos and gramtools use similar models – the gramtools model was based on that of Minos, but was modified to handle nested genotyping.

At each variant site, the aim is to choose the correct, ie most likely, allele from a set of alleles *A*. We have the following information from gramtools:

1. A function *γ* : *P* (*A*) *→* ℕ (where *P* (*A*) is the power set of *A*), defined by *γ*(*X*) = the number of reads that map to all alleles in *X* (and map to no other alleles). Since gramtools only reports the combinations of alleles that it sees, we define *γ* by assuming that all elements of *P* (*A*) not reported by gramtools have zero reads. Note also that gramtools does exact matching only - mismatches between the read and graph are not allowed;

2. For each allele *a* belonging to *A*, the read depth at each position in *a*, where reads are allowed to be multiply mapped. This means that for each allele gramtools matches a read to, a per-base coverage counter of each allele’s matching bases is incremented. Thus if a read maps to the middle 100bp of two long alternative alleles, then a counter is incremented at each position of those 100bp in each allele.

Let *c* be the total coverage at a site, given by

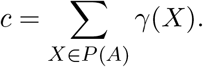

Let *a* be an allele belonging to *A*. Define the coverage *c*_*a*_ of *a* to be

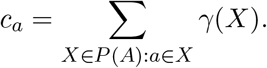

Let *ε* be the error rate in the reads, for which a default value of 0.002 is used and can be changed by the user. Let *d* be the expected read depth and *σ*^2^ the read depth variance, which are estimated using the read depth reported by gramtools at each variant site. We assume that the read depth follows a negative binomial distribution NB(*n, r*), where the parameters *n* and *r* are given by

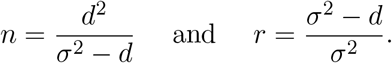

This requires *σ*^2^ *> d*. If this is not the case then we set *σ*^2^ to be double the read depth *d*. This is only expected to happen in rare circumstances and was only seen in the simulated data sets where the read depth was very even, unlike real data. The genotyping model used by Minos comprises the three terms:

1. “correct” coverage: NB(*n, r, c*_*a*_);

2. coverage due to read errors: 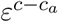;

3. a gap (ie zero coverage) penalty: 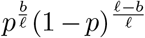, where *p* = 1 *−* NB(*n, r*, 0) is the probability that a given position has zero depth, *f* is the length of allele *a*, and *b* is the number of positions in *a* with non zero coverage.

The log likelihood is then calculated by summing the natural logarithm of these three terms. The allele with the greatest log likelihood is chosen, with genotype confidence of the difference in log likelihoods of that allele and the second greatest log likelihood. The genotype confidence is reported in the Minos output VCF file using the tag `GT_CONF`.

#### Variant call filtering

The FILTER column of the VCF file made by Minos is implemented using four filters. The first requires a read depth of at least two, called `MIN_DP` in the output VCF file. The second is a read depth no more than the mean depth plus three standard deviations, called `MAX_DP`. The third filter identifies apparent heterozygous calls (for example, caused by contamination), requiring by default at least 90% of the reads to support the called allele. It is called `MIN FRS` (“minimum fraction of read support”, can be set by the user). The final filter removes low confidence genotype calls. Since the genotyping model is dependent on read depth, the confidence score is not directly comparable between different sets of reads. This is accounted for by normalising the confidence score as follows. 10,000 SNPs are simulated by sampling read depths from a negative binomial distribution, defined by the observed mean depth and variance - this is the same distribution as used in the genotyping model. Incorrect read depth is sampled from a binomial distribution with *n* = observed read depth, and *p* = read error rate. These simulated SNPs are genotyped using the same method as used when variant calling, generating an expected distribution of genotyping confidence scores specific to this run. When genotyping the real variant sites, any call with a confidence score in the first 0.5% of the simulated genotype score distribution fails the filter (the threshold can be set by the user). This filter is called `MIN GCP` (minimum genotype confidence percentile) in the output VCF file.

#### Joint variant calling

The Minos Nextflow pipeline for joint genotyping large sets of samples is conceptually very similar to running on a per-sample basis, and proceeds as follows. The starting point is a VCF file of variant calls for each sample. These VCF files are clustered and merged, as described above, to produce a single gramtools graph for variant calling all samples. This graph should encapsulate all variants found across all of the input VCF files (except for deletions longer than 50bp). Each sample is genotyped using its reads mapped to the gramtools graph, resulting in a VCF file for each sample, where the variant sites are identical across all samples. The pipeline also produces a single multi-sample VCF file, combining the files using ivcfmerge (https://github.com/iqbal-lab-org/ivcfmerge). Finally, a distance matrix is calculated by defining the distance between any two samples to be the number of variant sites where those samples have different genotype calls.

### Variant call evaluation

The variant call evaluation with Varifier was implemented in Python and is available under the MIT license at https://github.com/iqbal-lab-org/varifier. The required input is: 1) a VCF file of variant calls to be evaluated; 2) a “mapping genome” FASTA file, which is the reference sequence corresponding to the VCF file; 3) a “truth genome”, which is the sequence assumed to be correct. The basic idea is to assign a score from zero (meaning false-positive) to one (true-positive) to each variant call, where fractional scores indicate partially correct calls. The score is determined by mapping probe sequences generated from the reference and alternative alleles to the truth genome.

### Precision

Each variant is processed using the following method. First, variants with no genotype call (`GT` field) or a heterozygous genotype are ignored. This method is designed for haploid organisms only since it essentially looks for perfect allele matches to the truth genome, which does not work for heterozygous genotype calls. A probe sequence is generated comprising the called allele, plus the (by default) 100 flanking nucleotides from the mapping reference before and after the allele - we call this the “alt probe”. Similarly, a “ref probe” is generated that uses the reference allele instead of the called allele. The alt probe is mapped to the truth genome using minimap2[42], and mappings that do not include the allele in the alignment or have mapping quality equal to zero are ignored. If there are no remaining mappings then the variant is classified as a false positive and assigned a score of zero. Otherwise, the minimap2 mapping that has the greatest number of allele positions matching the mapping genome is chosen as the “best” match. Similarly, the ref probe is mapped to the mapping genome and the best mapping is chosen, but with the additional requirement that the alignment start position in the mapping genome must be equal to that of the best alt probe mapping. If this results in no ref probe mapping, but with the alt probe mapped, then the variant is classified as a true positive and assigned a score of 1. If both probe sequences have a best match identified, then edit distances between sequences are used to define a score for the variant call. For motivation, consider the following relatively simple example allele sequences:

|  |  |
| --- | --- |
| Reference: | TAGAC |
| Alt allele: | TTGAA |
| Truth: | TTGAG |

Although the called allele is incorrect because it missed the C to G SNP, it does include the other A to T SNP and is 80% correct (4/5 of the sequence matches the truth). However, the called allele only contains half of the correct variation between the reference and the truth (1/2 SNPs are called), and we would like to account for this. Let *d*(*t, r*) be the edit distance between the truth and reference alleles, and *d*(*t, a*) the edit distance between the truth and alternative alleles. We define the score as:

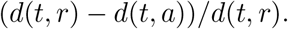

In the example above, the score is (2 *−* 1)*/*2 = 0.5. Note that in the simple case of a single SNP, a false positive scores zero and a true positive scores one. This edit distance-based measure is designed to handle indels and other complex variants. Although the score is usually between zero and one inclusive, in rare cases where the called allele is very distant from the truth, it is possible to have a negative score. The overall precision is calculated by dividing the total of the numerators by the total of the denominators, summed over all variants under consideration.

### Recall

Recall is determined using the following method (a flow chart is provided in Supplementary Figure 8). First, a VCF file of all expected calls must either be supplied by the user, or alternatively is made by comparing the mapping genome to the truth genome. Two separate expected callsets are made: one using MUMmer[43] and the other using minimap2 and PAFtools. MUMmer is used by running the commands from its `dnadiff` pipeline. minimap2 is run with the options `-c --cs`, and the output is piped into unix `sort -k6,6 -k8,8n`, and then into the `call` command of PAFtools with the options `-l50 -L50`. Taking the union gives a set of variant calls between the mapping and truth genomes, which we expect to contain false positives that need to be removed. The final truth variant callset is made using the same probe mapping method described in the above precision section, to remove false positive calls. Each variant call from MUMmer and minimap2 is kept if probe mapping to the truth genome results in a true positive call where the called allele matches perfectly (in other words, the variant has a score of one). In the rare case where both tools call at the same position but with conflicting calls, the calls are not used.

To recap, at this point in the recall pipeline we have a set of variants to be evaluated, a set of truth variant calls, and the mapping and truth genomes. The truth calls are in a VCF file with respect to the mapping genome. Next, all variants in the VCF file of calls to evaluate are applied to the mapping genome, to produce a new “mutated” genome. Determining recall is now the same as answering: how many truth variants are found in the mutated genome? This is answered using probe mapping using the same method as for precision. The VCF file of truth calls is evaluated, where the mutated reference takes the place of the “truth” genome.

## Benchmarking

### Truth genomes

The *S. aureus* truth genomes were generated from PacBio and Illumina reads as follows. The PacBio raw reads were assembled using canu v1.6[44] to produce the draft assembly. The contigs from the draft assembly were aligned to the respective reference genomes using the `nucmer` utility from the MUMmer3[45] package. The contigs were oriented to match the reference and trimmed based on the `nucmer` alignments and circularized using minimus2[46]. The assemblies were polished using the Illumina reads by iteratively running Pilon v1.23[47] until no more corections were made, up to a maximum of 10 runs. Reads were mapped using BWA MEM version 0.7.17 to make input for each Pilon iteration.

The 17 *M. tuberculosis* truth genomes were from [18], which already had the same Pilon polishing process applied to them as used on the *S. aureus* genomes. We used *K. pneumoniae* assemblies from [48] for the truth genomes, which we note already had Pilon run on them as part of their assembly process.

### Genome masks

A genome mask was available for *M. tuberculosis* H37Rv, which is used routinely by Public Health England. The plasmids were masked from the *K. pneumoniae* and *S. aureus* mapping genomes. A mask was generated for each truth genome by excluding positions where there was not a majority agreement between Illumina mapped Illumina reads and that genome, as described in the Supplementary Material. These masks were used by Varifier, which ignores all variants intersecting any masked region of the the truth or mapping genomes.

### *K. pneumoniae* reference genomes

The five *K. pneumoniae* genomes used as references for variant calling were chosen as follows. The average nucleotide identity (ANI) was calculated between each truth genome and all *K. pneumoniae* genomes in RefSeq using FastANI[49]. RefSeq[50] genomes with a minimum ANI less than 97.5% or a maximum ANI of 100% were excluded. The remaining genomes were listed in order of the minimum ANI, and 5 the genomes were chosen evenly spaced from this list, to obtain a range of ANI between the truth and mapping genomes.

### Joint genotyping

The nextflow pipeline included in the Minos github repository was used to run joint genotyping on all three data sets. A nextflow configuration file is included that contains preset profiles to set sensible default parameters for “medium” and “large” size data sets. The “medium” profile was used for the Walker 2013 data set, and “large” was used for the CRyPTIC and Mykrobe data. The pipeline needs an input TSV file, listing sample names and paths to VCF and reads files, plus the reference genome in FASTA format, and optionally a reference genome mask in BED format. The single command line used to run the complete pipeline is provided in the Supplementary Material. The pipeline is set up to handle large cohorts by saving RAM where possible. It splits the genome into slices, each containing approximately the same number of alleles, and processes each slice in series. Each slice overlaps by a read length to remove end effects from mapping. The Walker 2013 set was split into 100 slices, and the two large sets into 300 slices. The slicing option requires that the input reads for each sample are in a sorted, indexed BAM file, so that the reads for each slice can be efficiently extracted. Such BAM files are typically generated during most variant calling pipelines, and so this requirement is unlikely to create further work.

Joint genotyping with BayesTyper and GraphTyper both required a single sorted VCF file from concatenating (taking account of VCF headers) all input VCF files. This file was sorted, compressed with bgzip, and indexed with tabix for GraphTyper. For BayesTyper, bcftools norm was run on the VCF file. Then the same commands used for running on individual samples were used, as described in the Supplementary Material. These were successful on all samples in the Walker 2013 data set. On the Mykrobe data set both tools failed on the first sample, ERR025833. For BayesTyper, the `combine` function ran successfully, but the `cluster` command failed. GraphTyper failed when running the `genotype` function, after processing approximately 10% of the genome, with a peak RAM usage of 76GB.

After joint genotyping the CRyPTIC and Mykrobe data sets, the RRDR region was analysed as follows. To avoid ambiguity, we note that the RRDR region is the 27 amino acid sequence at 426-452 in the *rpoB* gene in the H37Rv genome (it is often alternatively described with *E. coli* numbering as 507-533). In H37Rv genome coordinates this is 759807-763325.

To analyse the RRDR region, only samples that had a high quality rifampicin phenotype of resistant or susceptible were used. This information is provided in the supplementary tables of [29] for the Mykrobe data. For the CRyPTIC data, we used the samples from [10]. Each of these samples was processed as follows. Its variants contained in the whole *rpoB* gene were extracted and applied to the genome sequence and translated, making a mutated amino acid sequence. The amino acid variants of the sample were deduced from aligning the mutated amino acid sequence to the reference amino acid sequence. Then the variants in the RRDR were extracted for each sample. In this way, combinations of nucleotide variants were accounted for (for example, two consecutive SNPs could cause a single amino acid change).

## Data Availability

The reference genomes used for variant calling were as follows.

- *M. tuberculosis* : H37Rv version 3 reference genome NC_000962.3[23].
- *S. aureus*: USA300 genome GCA_000013465.1[51] and the TW20 genome GCA_000027045.1[52].
- *K. pneumoniae*: GCF_000784945[53], GCF_001952915[54], GCF_003073315, GCF_003076555[55], and GCF_011006575[56].

A data download is available at https://doi.org/10.6084/m9.figshare.16613398 that contains FASTA files of all truth genomes and BED files of all genome masks. Accessions for reads and assemblies are provided in Supplementary Tables 8–10, and are also included in the data download in tab-delimited format. The Mykrobe data set accessions are in the Supplementary file from [29], available at https://doi.org/10.6084/m9.figshare.7556789. Details of the CRyPTIC data can be found in [10].

The pipeline to call variants, run BayesTyper, GraphTyper, and Minos, and run Varifier was written in Python and is available under the MIT license at https://github.com/iqbal-lab-org/minos-paper-benchmarking. A copy of the Singularity[57] container used for this study is available at https://doi.org/10.6084/m9.figshare.16613383. All software versions and command lines used are in the Supplementary Material.

## Acknowledgements

The authors thank Robyn Ffrancon for software engineering in gramtools, and Sara Goodwin from Cold Spring Harbour for assistance with PacBio sequencing.

## Author Contributions

The study was conceived by ZI. MH developed Minos and Varifier and ran all benchmarking. BL developed gramtools functionality used by Minos, and contributed to the genotyping model and Minos code. GN wrote the ivcfmerge code, used by joint genotyping. MBH and LL helped design Varifier, and MBH wrote parts of the Varifier code. RC contributed to the genotyping model. MS and SR assembled and quality-checked the *S. aureus* assemblies. MH and ZI drafted the manuscript. KM helped with the RRDR analysis and edited the manuscript. All authors reviewed and approved the manuscript.

## CRyPTIC consortium

Members of the CRyPTIC consortium (in alphabetical order):

Ivan Barilar^1^, Simone Battaglia^2^, Emanuele Borroni^2^, Angela Pires Brandao^3,4^, Alice Brankin^5^, Andrea Maurizio Cabibbe^2^, Joshua Carter^6^, Daniela Maria Cirillo^2^, Pauline Claxton^7^, David A Clifton^5^, Ted Cohen^8^, Jorge Coronel^9^, Derrick W Crook^5^, Viola Dreyer^1^, Sarah G Earle^5^, Vincent Escuyer^10^, Lucilaine Ferrazoli^4^, Philip W Fowler^5^, George Fu Gao^11^, Jennifer Gardy^12^, Saheer Gharbia^13^, Kelen Teixeira Ghisi^4^, Arash Ghodousi^2,14^, Ana Luíza Gibertoni Cruz^5^, Louis Grandjean^15^, Clara Grazian^16^, Ramona Groenheit^17^, Jennifer L Guthrie^18,19^, Wencong He^11^, Harald Hoffmann^20,21^, Sarah J Hoosdally^5^, Martin Hunt^22,5^, Zamin Iqbal^22^, Nazir Ahmed Ismail^23^, Lisa Jarrett^24^, Lavania Joseph^23^, Ruwen Jou^25^, Priti Kambli^26^, Rukhsar Khot^26^, Jeff Knaggs^22,5^, Anastasia Koch^27^, Donna Kohlerschmidt^10^, Samaneh Kouchaki^5,28^, Alexander S Lachapelle^5^, Ajit Lalvani^29^, Simon Grandjean Lapierre^30^, Ian F Laurenson^7^, Brice Letcher^22^, Wan-Hsuan Lin^25^, Chunfa Liu^11^, Dongxin Liu^11^, Kerri M Malone^22^, Ayan Mandal^31^, Mikael Mansjö^17^, Daniela Matias^24^, Graeme Meintjes^27^, Flávia de Freitas Mendes^4^, Matthias Merker^1^, Marina Mihalic^21^, James Millard^32^, Paolo Miotto^2^, Nerges Mistry^31^, David Moore^33,9^, Kimberlee A Musser^10^, Dumisani Ngcamu^23^, Hoang Ngoc Nhung^34^, Stefan Niemann^1,35^, Kayzad Soli Nilgiriwala^31^, Camus Nimmo^15^, Nana Okozi^23^, Rosangela Siqueira Oliveira^4^, Shaheed Vally Omar^23^, Nicholas Paton^36^, Timothy EA Peto^5^, Juliana Maira Watanabe Pinhata^4^, Sara Plesnik^21^, Zully M Puyen^37^, Marie Sylvianne Rabodoarivelo^38^, Niaina Rakotosamimanana^38^, Paola MV Rancoita^14^, Priti Rathod^24^, Esther Robinson^24^, Gillian Rodger^5^, Camilla Rodrigues^26^, Timothy C Rodwell^39,40^, Aysha Roohi^5^, David Santos-Lazaro^37^, Sanchi Shah^31^, Thomas Andreas Kohl^1^, Grace Smith^24,13^, Walter Solano^9^, Andrea Spitaleri^2,14^, Philip Supply^41^, Utkarsha Surve^26^, Sabira Tahseen^42^, Nguyen Thuy Thuong Thuong^34^, Guy Thwaites^34,5^, Katharina Todt^21^, Alberto Trovato^2^, Christian Utpatel^1^, Annelies Van Rie^43^, Srinivasan Vijay^44^, Timothy M Walker^5,34^, A Sarah Walker^5^, Robin Warren^45^, Jim Werngren^17^, Maria Wijkander^17^, Robert J Wilkinson^46,47,29^, Daniel J Wilson^5^, Penelope Wintringer^22^, Yu-Xin Xiao^25^, Yang Yang^5^, Zhao Yanlin^11^, Shen-Yuan Yao^23^, Baoli Zhu^48^.

^1^Research Center Borstel, Borstel, Germany

^2^IRCCS San Raffaele Scientific Institute, Milan, Italy

^3^Oswaldo Cruz Foundation, Rio de Janeiro, Brazil

^4^Institute Adolfo Lutz, São Paulo, Brazil

^5^University of Oxford, Oxford, UK

^6^Stanford University School of Medicine, Stanford, USA

^7^Scottish Mycobacteria Reference Laboratory, Edinburgh, UK

^8^Yale School of Public Health, Yale, USA

^9^Universidad Peruana Cayetano Heredia, Lima, Perú

^10^Wadsworth Center, New York State Department of Health, Albany, USA

^11^Chinese Center for Disease Control and Prevention, Beijing, China

^12^Bill & Melinda Gates Foundation, Seattle, USA

^13^UK Health Security Agency, London, UK

^14^Vita-Salute San Raffaele University, Milan, Italy

^15^University College London, London, UK

^16^University of New South Wales, Sydney, Australia

^17^Public Health Agency of Sweden, Solna, Sweden

^18^The University of British Columbia, Vancouver, Canada

^19^Public Health Ontario, Toronto, Canada

^20^SYNLAB Gauting, Munich, Germany

^21^Institute of Microbiology and Laboratory Medicine, IMLred, WHO-SRL Gauting, Germany

^22^EMBL-EBI, Hinxton, UK

^23^National Institute for Communicable Diseases, Johannesburg, South Africa

^24^Public Health England, Birmingham, UK

^25^Taiwan Centers for Disease Control, Taipei, Taiwan

^26^Hinduja Hospital, Mumbai, India

^27^University of Cape Town, Cape Town, South Africa

^28^University of Surrey, Guildford, UK

^29^Imperial College, London, UK

^30^Université de Montréal, Canada

^31^The Foundation for Medical Research, Mumbai, India

^32^Africa Health Research Institute, Durban, South Africa

^33^London School of Hygiene and Tropical Medicine, London, UK

^34^Oxford University Clinical Research Unit, Ho Chi Minh City, Viet Nam

^35^German Center for Infection Research (DZIF), Hamburg-Lübeck-Borstel-Riems, Germany

^36^National University of Singapore, Singapore

^37^Instituto Nacional de Salud, Lima, Perú

^38^Institut Pasteur de Madagascar, Antananarivo, Madagascar

^39^FIND, Geneva, Switzerland

^40^University of California, San Diego, USA

^41^Univ. Lille, CNRS, Inserm, CHU Lille, Institut Pasteur de Lille, U1019 - UMR 9017 - CIIL - Center for Infection and Immunity of Lille, F-59000 Lille, France

^42^National TB Reference Laboratory, National TB Control Program, Islamabad, Pakistan

^43^University of Antwerp, Antwerp, Belgium

^44^University of Edinburgh, Edinburgh, UK

^45^Stellenbosch University, Cape Town, South Africa

^46^Wellcome Centre for Infectious Diseases Research in Africa, Cape Town, South Africa

^47^Francis Crick Institute, London, UK

^48^Institute of Microbiology, Chinese Academy of Sciences, Beijing, China

CRyPTIC consortium competing interests:

E.R. is employed by Public Health England and holds an honorary contract with Imperial College London. I.F.L. is Director of the Scottish Mycobacteria Reference Laboratory. S.N. receives funding from German Center for Infection Research, Excellenz Cluster Precision Medicine in Chronic Inflammation, Leibniz Science Campus Evolutionary Medicine of the LUNG (EvoL-UNG)tion EXC 2167. P.S. is a consultant at Genoscreen. T.R. is funded by NIH and DoD and receives salary support from the non-profit organization FIND. T.R. is a co-founder, board member and shareholder of Verus Diagnostics Inc, a company that was founded with the intent of developing diagnostic assays. Verus Diagnostics was not involved in any way with data collection, analysis or publication of the results. T.R. has not received any financial support from Verus Diagnostics. UCSD Conflict of Interest office has reviewed and approved T.R.’s role in Verus Diagnostics Inc. T.R. is a co-inventor of a provisional patent for a TB diagnostic assay (provisional patent #: 63/048.989). T.R. is a co-inventor on a patent associated with the processing of TB sequencing data (European Patent Application No. 14840432.0 & USSN 14/912,918). T.R. has agreed to “donate all present and future interest in and rights to royalties from this patent” to UCSD to ensure that he does not receive any financial benefits from this patent. S.S. is working and holding ESOPs at HaystackAnalytics Pvt. Ltd. (Product: Using whole genome sequencing for drug susceptibility testing for Mycobacterium tuberculosis). G.F.G. is listed as an inventor on patent applications for RBD-dimer-based CoV vaccines. The patents for RBD-dimers as protein subunit vaccines for SARS-CoV-2 have been licensed to Anhui Zhifei Longcom Biopharmaceutical Co. Ltd, China.

