## Supplementary Material for "Minos: variant adjudication and joint genotyping of cohorts of bacterial genomes"

[1]European Bioinformatics Institute, Cambridge, UK
[2]Nuffield Department of Medicine, University of Oxford, Oxford, UK
[3]Institute of Evolutionary Biology, Ashworth Laboratories, University of Edinburgh, UK
[4]Department of Computer Science, Johns Hopkins University, Baltimore, MD, USA
*Please see main manuscript section "CRyPTIC consortium" for details.

#### Contents

**1 Simulated data** 2

**2 Conflicting Joint Genotyping Calls** 2

**3 Software versions and command lines** 3

**4 Figures** 7

**5 Tables** 14

### 1 Simulated data

Simulations were performed as a sanity check on performance of all benchmarked tools, and were not intended (and are not claimed) to be good representatives of real-world data. They do however provide a way to potentially reveal limitations of tools. Simulated data sets were generated from the *M. tuberculosis* genome H37Rv using simutator (`https://github.com/iqbal-lab-org/simutator`). A mutated genome was generated for each type of variation, with variants distributed uniformly across the genome: a SNP every 200bp, an insertion of length 1 every kilobase, a deletion of length 1 every kilobase etc. Insertions and deletions were simulated up to a length of 10kbp. Paired Illumina reads were simulated at 50X depth from each mutated reference sequence using ART.

BayesTyper, GraphTyper, and Minos achieved near-perfect precision and recall on all data sets, showing that all tools are effective on clean data (Supplementary Tables 1, 2). All three tools had perfect precision when calling isolated SNPs, insertions and deletions. They also had perfect recall of SNPs, except for Minos with a recall of 99.68% using its default call filters. However, including all Minos calls results in perfect recall (and still perfect precision). This trend continued in the remainder of the simulation results. We found that the Minos default filters are negatively affected by the very clean simulated reads with idealised read depth distributions. As shown later, the filters are however effective on real data. The recall of SNPs and the full range of indels tested is shown in Supplementary figure 1. Again, all tools have near 100% recall, except filtered Minos calls which loses up to 0.9% recall on small ($< 50$bp) variants.

In addition to isolated SNPs, insertions, and deletions, four data sets were generated with a complex variants every kilobase, with each complex variant consisting of SNPs and/or indels of varying length in a window of size 10 to 50bp. The results were very similar across all tools (Supplementary Table 2).

### 2 Conflicting Joint Genotyping Calls

As noted in the main text, Minos resolves overlapping input variants, so that no two sites in its output contain reference positions in common. Furthermore, this means Minos cannot output two separate VCF records with incompatible genotype calls, in contrast to BayesTyper and GraphTyper. A typical example is shown next.

On the Walker 2013 outbreak data set, BayesTyper and GraphTyper reported the following two lines in their output VCF files for each sample (first five columns shown only for brevity – these are enough to describe the variants that are being called):

```
NC_000962.3     397272  .       TCGGCGCC        T
NC_000962.3     397275  .       G               C,*
```

BayesTyper and GraphTyper made inconsistent genotype calls of these variants. For example in sample ERR046906, BayesTyper and GraphTyper genotyped the deletion at 397272 as not present (genotype `0` or `0/0`), saying in particular that this sample has a `G` at position 397275. However, both tools also genotyped the second variant line at 397275 as having a `C` (genotype `1` or `1/1`). These genotype calls cannot both be correct.

Minos reported the SNP and deletion alleles with the following single line in its VCF output file:

```
NC_000962.3     397272  .       TCGGCGCC        T,TCGCCGCC
```

and called the second alternative allele `TCGCCGCC` as correct – this is a SNP from `G` to `C` at position 397275. Manual inspection of the reads pileup confirmed that this is the correct call.

#### 3 Software versions and command lines

All software was run using a Singularity container, which was made from the definition file `singularity.def` in the github repository `https://github.com/iqbal-lab-org/minos-paper-benchmarking` (git commit dbbc62a9388fd30869406cc63c10e70c08d45a0b).

##### Simulated data

The simulated genomes were made using Simutator (`https://github.com/iqbal-lab-org/simutator`) git commit 0218c8a5b37fd72eb4e5b2df4cba9f6118f96788. The command line was:

```
simutator mutate_fasta --seed 42 --snps 200 \
  --dels 1000:1,1000:2,1000:3,1000:4,1000:5,1000:10,1000:20,1000:50,10000:100,\
20000:500,20000:1000,20000:2000,20000:5000,20000:10000 \
  --ins 1000:1,1000:2,1000:3,1000:4,1000:5,1000:10,1000:20,1000:50,10000:100,
20000:500,20000:1000,20000:2000,20000:5000,20000:10000 \
  --complex 1000:10:3:0:0:0,1000:10:3:1:1:2,1000:50:10:1:1:3,1000:50:20:3:3:4 \
  tb_ref.h37rv.fa out
```

ART version 20160605 (`https://www.niehs.nih.gov/research/resources/assets/docs/artbinmountrainier20160605linux64tgz.tgz`) was used to simulate Illumina reads from each of the mutated genomes output by simutator, using the command:

```
art_illumina --in mutated.fa --out reads.out \
  --noALN --seqSys HS25 --len 150 --fcov 50 \
  --mflen 500 --sdev 25 --rndSeed 42
```

##### Variant calling pipeline

The pipeline described next is implemented in the `minos-paper-benchmarking` container as a single script that runs all stages on one sample. First, all the relevant genome index files must be made (for example BWA index). This needs to be run once on each mapping reference FASTA file:

```
clockwork reference_prepare --outdir ref_dir mapping_reference.fasta
```

to make a directory (in this example, called `ref_dir`), which is used by the next command. The whole variant calling and evaluation pipeline was then run on one sample with the command

```
minospb run_one_sample \
  --truth_mask_bed truth_mask.bed \
  --ref_mask_bed ref_mask.bed \
  sample_name  out_directory ref_dir \
  reads_1.fastq.gz reads_2.fastq.gz
```

The stages run by that script are described next. First, reads were trimmed using Trimmomatic version 0.36, with the options

```
ILLUMINACLIP:/Trimmomatic-0.36/adapters/TruSeq3-PE-2.fa:2:30:10
LEADING:10 TRAILING:10 SLIDINGWINDOW:4:15 MINLEN:50 -phred33
```

where `/Trimmomatic/` is the root directory of the Trimmomatic download.

Reads were mapped with BWA MEM version 0.7.17 with option `-M`, then The output SAM file was sorted using SAMtools version 1.10.2 `samtools sort`, and then duplicates removed with `samtools rmdup` (defaults used for both commands), to make a file called `rmdup.bam`.

Variant calls were made with SAMtools and BCFtools version 1.10.2 with the command:

```
samtools mpileup -ugf rmdup.bam | bcftools call -vm -O v -o out.vcf
```

Cortex git commit 3a235272e4e0121be64527f01e73f9e066d378d3 (and dependencies VCFtools version 0.1.15 and Stampy version 1.0.32) was used with the Cortex Perl script to call variants:

```
run_calls.pl --fastaq_index in.index --auto_cleaning yes --first_kmer 31 \
  --bc yes --pd no --outdir cortex.out --outvcf cortex --ploidy 2 \
  --stampy_hash ref.stampy --stampy_bin /path/to/stampy.py \
  --list_ref_fasta /path/to/refs_file --refbindir /path/to/ref_dir \
  --genome_size N --qthresh 5 --mem_height 22 --mem_width 100 \
  --vcftools_dir /path/to/vcftools --do_union yes --ref CoordinatesAndInCalling \
  --workflow independent --logfile out.log
```

##### BayesTyper

BayesTyper version 1.5 was used, with the dependency KMC version 3.1.1. KMC was run with the command

```
kmc -t1 -m7 -k55 -ci1 -fbam rmdup.bam 01.kmc .
```

followed by

```
bayesTyperTools makeBloom -k 01.kmc
```

Then for each of the SAMtools and Cortex VCF files:

```
bcftools norm -f /path/to/ref.fa <samtools|cortex>.vcf \
  -o 02.typertools.<samtools|cortex>.norm.vcf
```

Then the commands:

```
bayesTyperTools combine -v samtools:samtools.norm.vcf,cortex:cortex.norm.vcf \
  -o 02.typertools.combine -z

bayesTyper cluster -r 42 -v 02.typertools.combine.vcf.gz -s \
  03.typertools.samples.tsv -g /path/to/ref.fa

bayesTyper genotype --noise-genotyping -r 42 \
  -v bayestyper_unit_1/variant_clusters.bin \
  -y 04.typertools.ploidy.tsv -c bayestyper_cluster_data \
  -s 03.typertools.samples.tsv -g /path/to/ref.fa -o bayestyper_unit_1/bayestyper
```

##### GraphTyper

GraphTyper version 2.5.1 was used, together with custom Python code to glue the various stages together and other required processing. These commands were used on the input VCF files:

```
bcftools concat -a -O v <VCF files> | bcftools sort -Oz -o concat.vcf.gz

tabix -p vcf concat.vcf.gz
```

to make a sorted bgzipped indexed VCF file.

Then to genotype using the default mode:

```
graphtyper genotype --output 01.genotype -vverbose --sam rmdup.bam \
  --region_file 00.regions.txt --threads 1 /path/to/ref.fa \
  --vcf concat.vcf.gz

bcftools concat -a -O v <list of all VCF files made by previous command>
```

Alternatively, to genotype using SV mode:

```
graphtyper genotype_sv --output 01.genotype -vverbose --sam rmdup.bam \
  --region_file 00.regions.txt --threads 1 /path/to/ref.fa concat.sv.vcf.gz

bcftools concat -a -O v <list of all VCF files made by previous command>
```

#### Minos

Minos git commit c22f4b00c61be7854336dad5b31b44c0d11e8e81 was used, and dependencies gramtools (git commit 8af53f6c8c0d72ef95223e89ab82119b717044f2), vcflib (git commit da9929c00938b1d1b042e8ce42514211ef34875d), and vt (git commit a549707d8e33006ed89c46ca7fbda007123418ab). The command line was:

```
minos adjudicate --reads rmdup.bam outdir /path/to/ref.fa samtools.vcf cortex.vcf
```

#### Genome masks

An existing genome mask was used for the *M. tuberculosis* H37Rv reference genome, which is routinely used by Public Health England variant calling pipeline COMPASS.

For all other genomes, masks for each mapping and truth genome were generated using agreement between the reads and the genomes as follows. The whole process is wrapped in a single script, run with the command:

```
minospb make_mask genome.fasta reads.1.fastq.gz reads.2.fastq.gz outdir
```

Illumina reads expected to perfectly match the genome (ie reads that made the genome assembly sequence) were trimmed, mapped, and PCR duplicates removed using exactly the same method as described earlier for variant calling. The resulting BAM file is parsed using pysam. A position is included in the mask if the depth of reads matching the reference is less than 5, or if the percent of reads that match the reference is less than 90%. The output is a mask in BED file format, which was then used as input to Varifier (see next section).

#### Varifier

varifier git commit ad3d6db3b851eb46357b0308e7496580e191700f was used, with the command line

```
varifier vcf_eval  --ref_mask ref_mask.bed \
  --truth_mask truth_mask.bed --filter_pass PASS \
  truth_ref.fasta mapping_ref.fasta to_evaluate.vcf outdir
```

To evaluate all calls made by a tool in order to see the effect of the FILTER column in the VCF, the option `--filter_pass` was omitted.

##### Joint Genotyping

The per-sample VCF files were generated using the variant calling method described above, making calls from SAMtools and Cortex. For each sample, these were input into Minos to make a single VCF for each sample. This was run using the script described above `minospb run_one_sample`.

These per-sample VCF files were used as input to the Minos joint genotyping pipeline, which is implemented using Nextflow. The Nextflow script and configuration file are in the `nextflow/` directory of the Minos repository, called `regnotype.nf` and `regenotype.config`. On the Walker 2013 data set, the pipeline was run with the command:

```
nextflow run \
  -w /path/to/working/dir/
  -with-singularity /path/to/minospg_image \
  -c regenotype.config \
  -profile medium \
  regnotype.nf
  --ref_fasta /path/to/tb_ref.h37rv.fa \
  --make_distance_matrix \
  --manifest manifest.tsv \
  --mask_bed_file tb.h37rv.mask.R00000039_repregions.bed \
  --outdir output_directory
```

where `manifest.tsv` is the required manifest file, containing for each sample its name, input VCF file, and BAM file of reads. The reads must be in a mapped, sorted, indexed BAM file for the pipeline. This enables it to split the reference into chunks and map reads to each chunk, which saves run time and memory. On the large CRyPTIC and Mykrobe data sets, the option `-profile large` was used, which allocates more memory and CPUs to tasks where necessary, and the option `--make_distance_matrix` was omitted.

#### 4 Figures

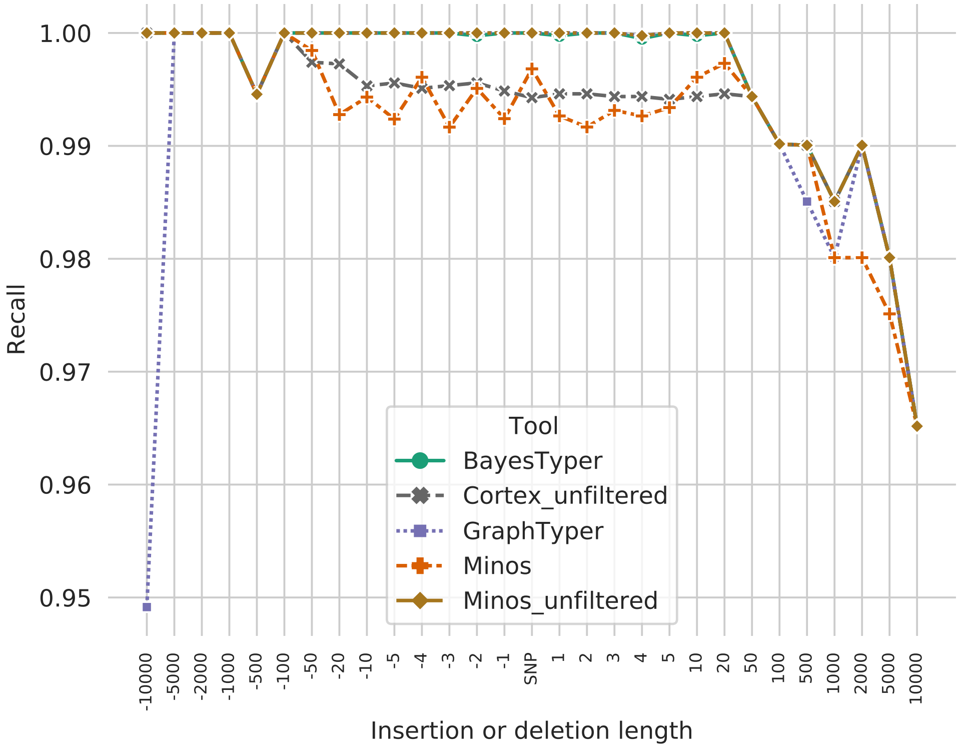


**Supplementary Figure 1:** Recall of isolated deletions, SNPs, and insertions on the simulated data set. Negative length values show deletions, and positive values show insertions. SNPs are shown in the centre. Note that the $x$-axis is non-linear.

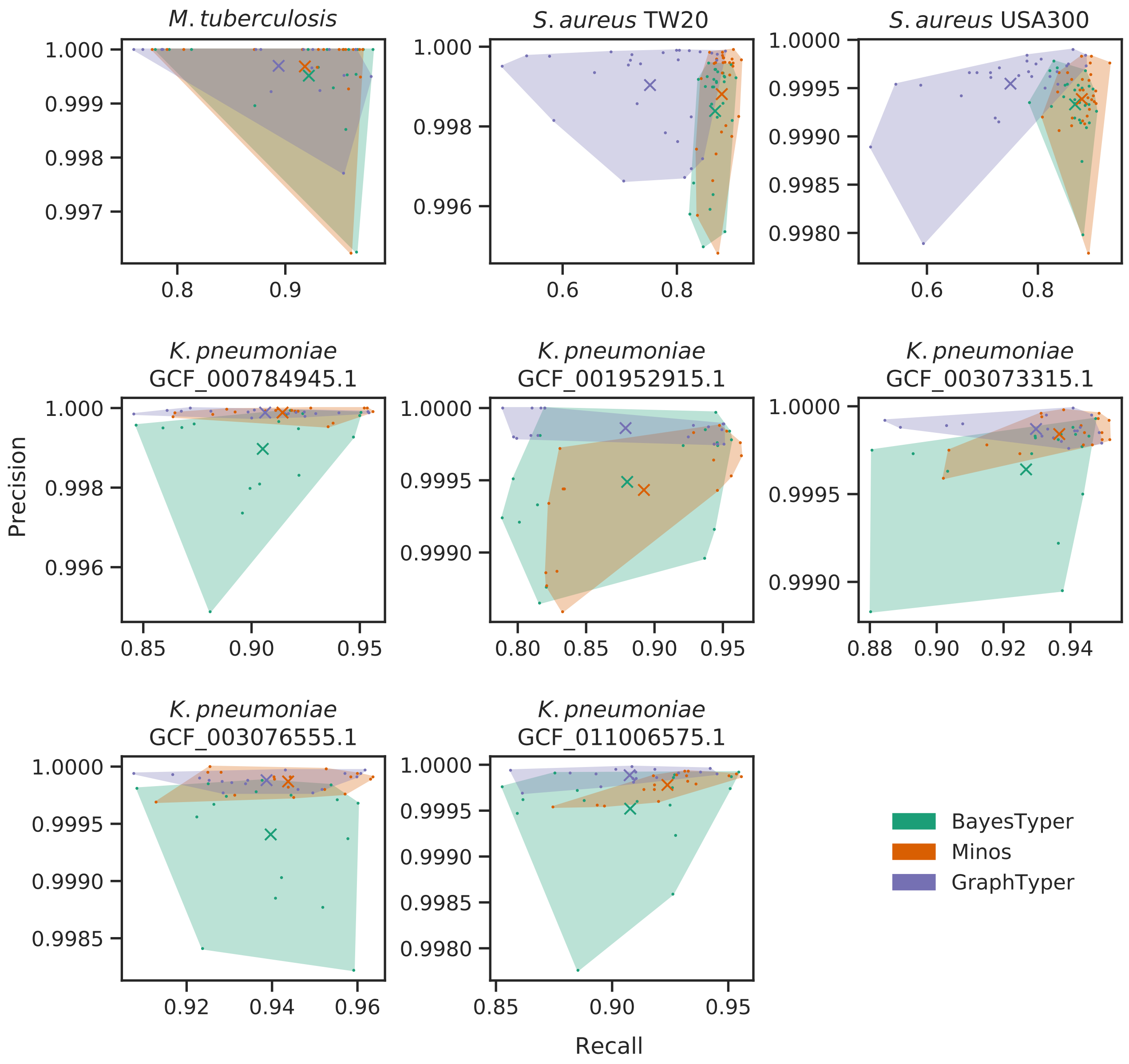


**Supplementary Figure 2:** Comparison of BayesTyper, GraphTyper and Minos on the bacteria data set. Precision and recall of each sample is shown as a dot, and the mean precision and recall for each tool is marked with a cross. The convex hull of the data points for each caller is shaded with an associated colour. Note the axis scales are different between the plots.

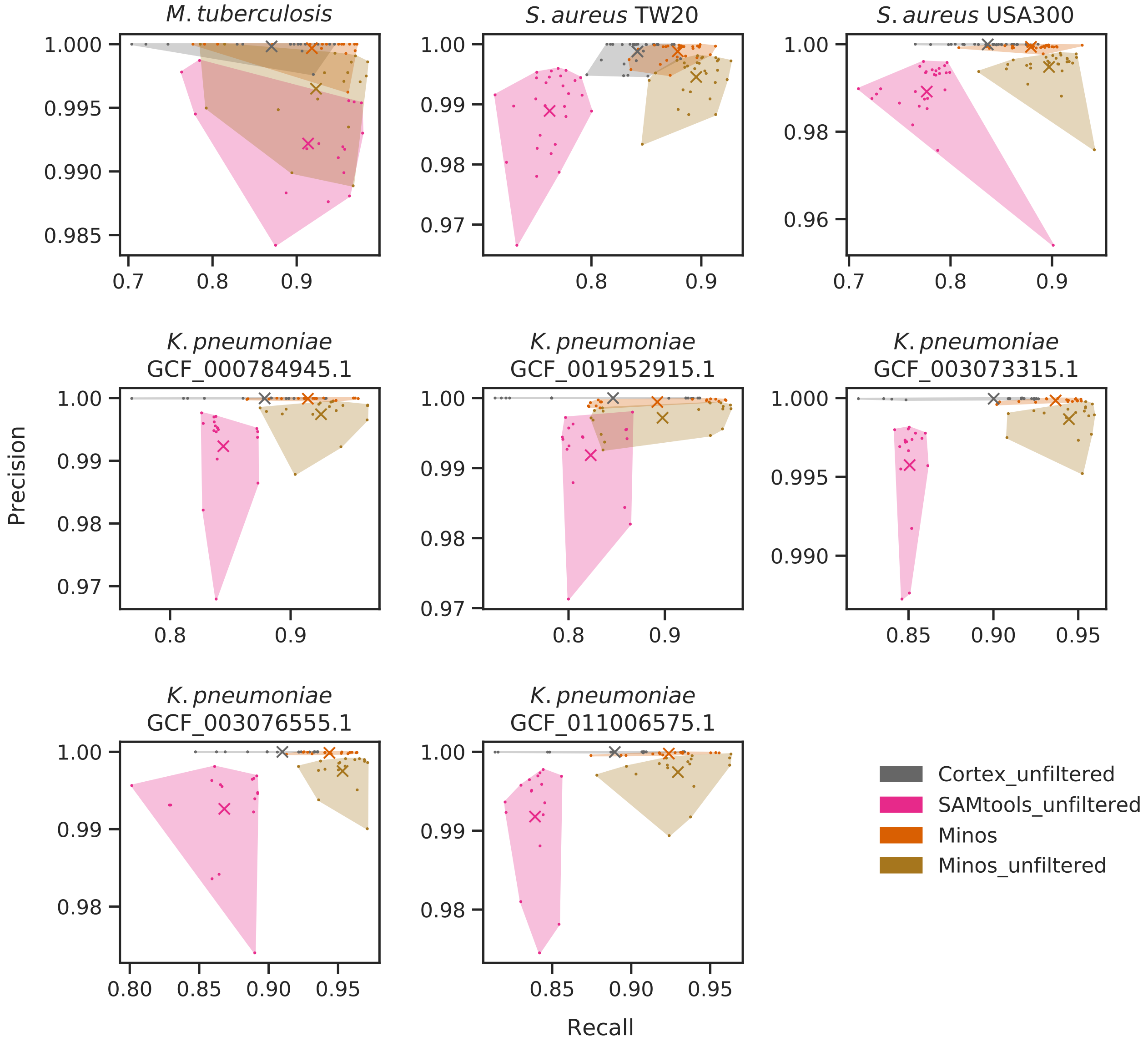


**Supplementary Figure 3:** Comparison of filtered and unfiltered Minos results, together with unfiltered Cortex and unfiltered SAMtools on the bacteria data set. Precision and recall of each sample is shown as a dot, and the mean precision and recall for each tool is marked with a cross. The convex hull of the data points for each caller is shaded with an associated colour. Note the axis scales are different between the plots.

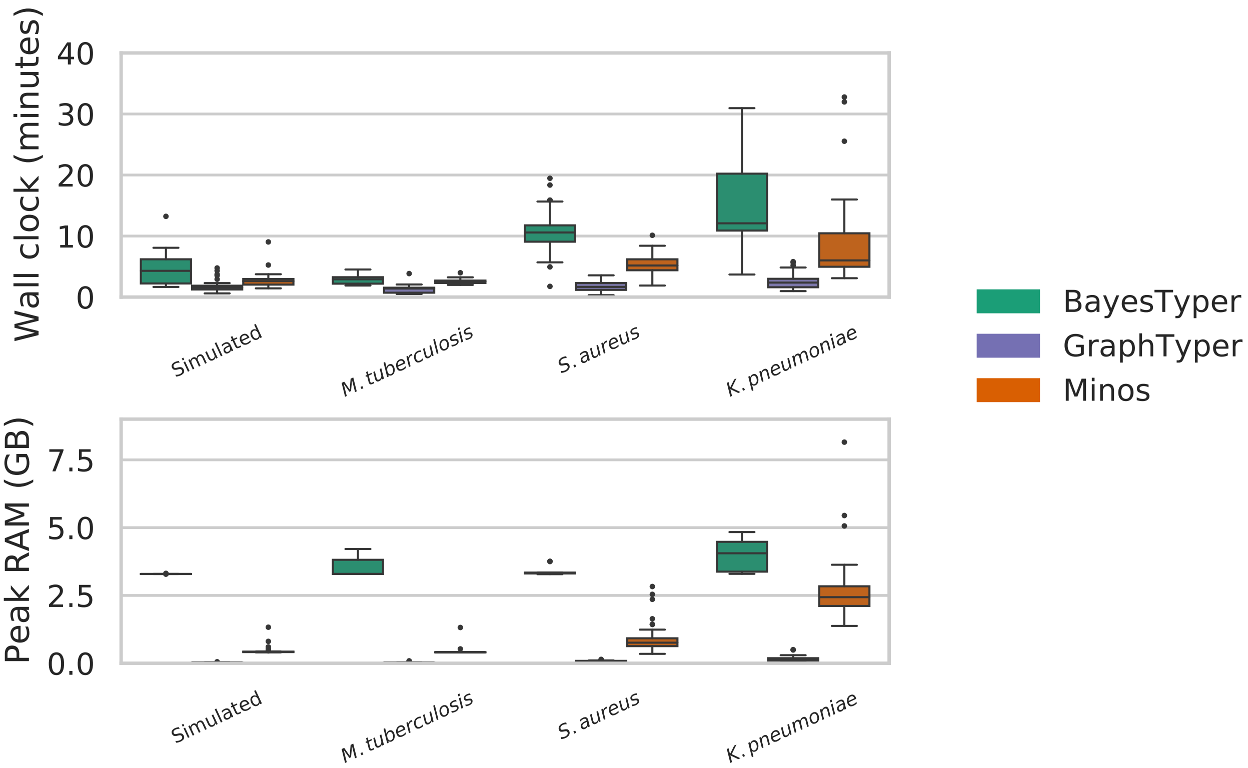


**Supplementary Figure 4:** Run time and peak RAM usage on the simulated and bacteria data sets. Values are taken from the output of the Unix command `time -v`.

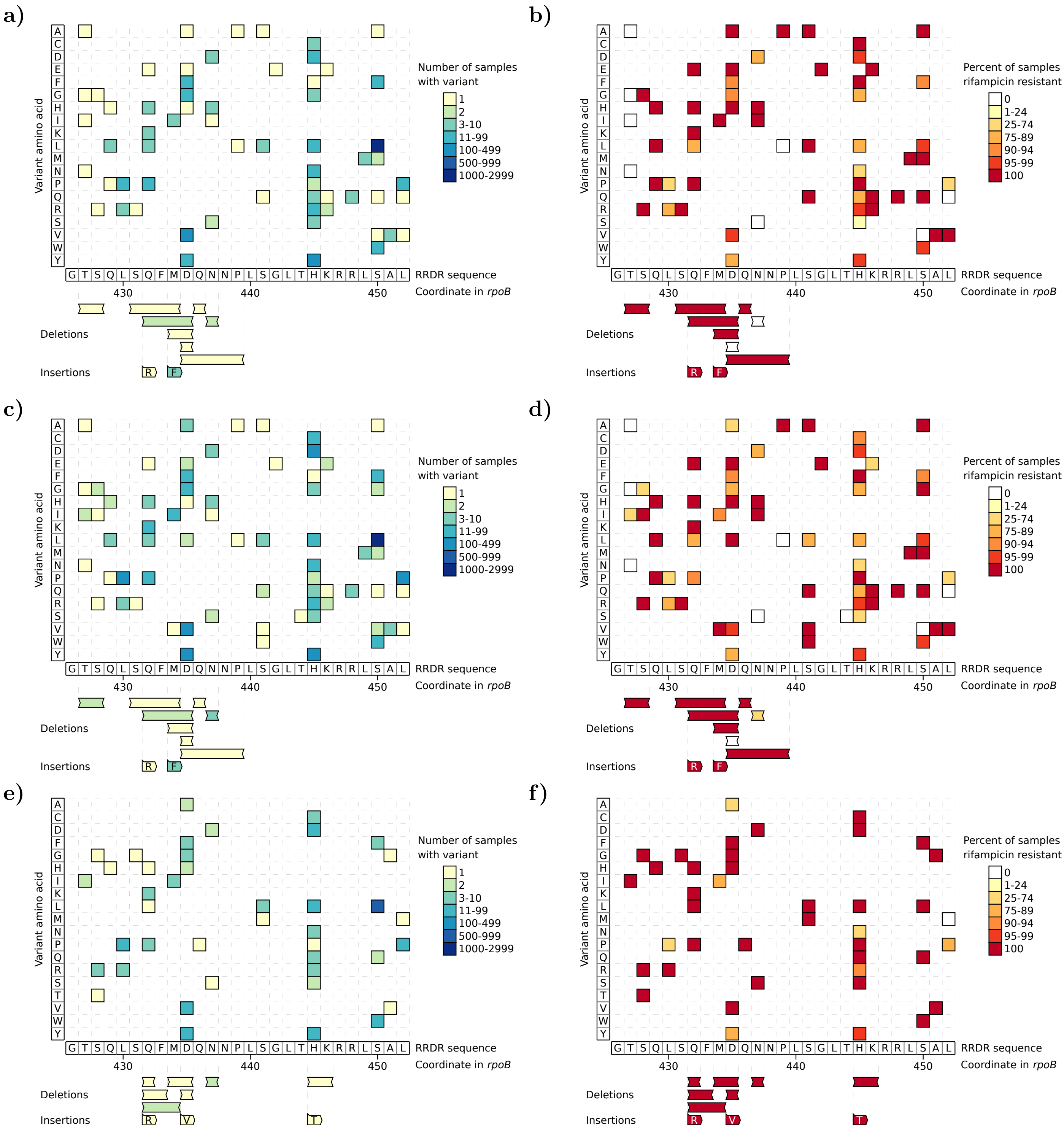


**Supplementary Figure 5:** All amino acid variants identified in the RRDR of the rpoB gene by joint genotyping the CRyPTIC and Mykrobe *M. tuberculosis* data sets. a) and b) are the same as Figure 4 in the main manuscript, showing samples from the CRyPTIC data set that have high quality phenotypes only. c) and d) again show the CRyPTIC set, but include all samples regardless of phenotype quality. e) and f) show the Mykrobe data set. In a), c), e) each variant is coloured by the number of samples possessing that variant. Plots b), d), f) colour the variants by the percent of samples with that variant that are rifampicin resistant. Each plot shows the RRDR region from left to right. Single amino acid variants are shown in the upper grid, with the y axis corresponding to the variant amino acid. The lower area shows deletions and insertions, with the inserted sequence given in the coloured boxes. For example, in a) and b) the leftmost deletion of amino acids TS at position 427-428 is found in one resistant sample. The leftmost insertion adds R after the S at position 431.

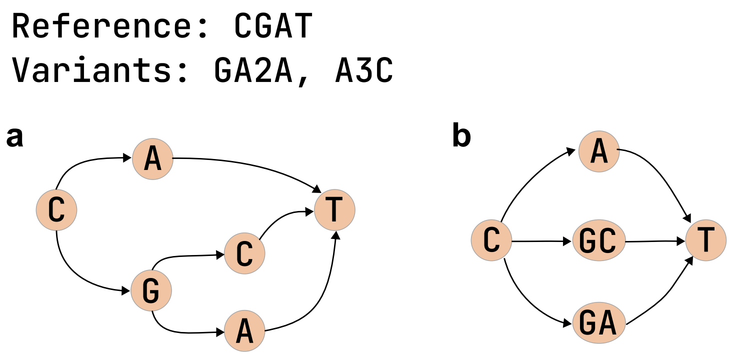


**Supplementary Figure 6:** a) Nested and b) non-nested graphs. The nested graph in a) is not allowed, and is transformed by Minos into b) for graph construction and then read mapping. Note that the two graphs contain the same sequences.

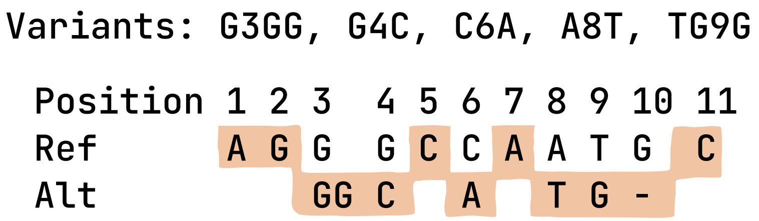


**Supplementary Figure 7:** Duplicated paths. The listed variants when applied to the reference sequence `AGGGCCAATGC` result in the possible reference and alternative alleles shown. The path marked in orange shows an allele combination that results in exactly the same sequence as the reference sequence. Minos prevents this situation from arising, so that reads cannot have the option of mapping to two different paths that are in fact the same sequence.

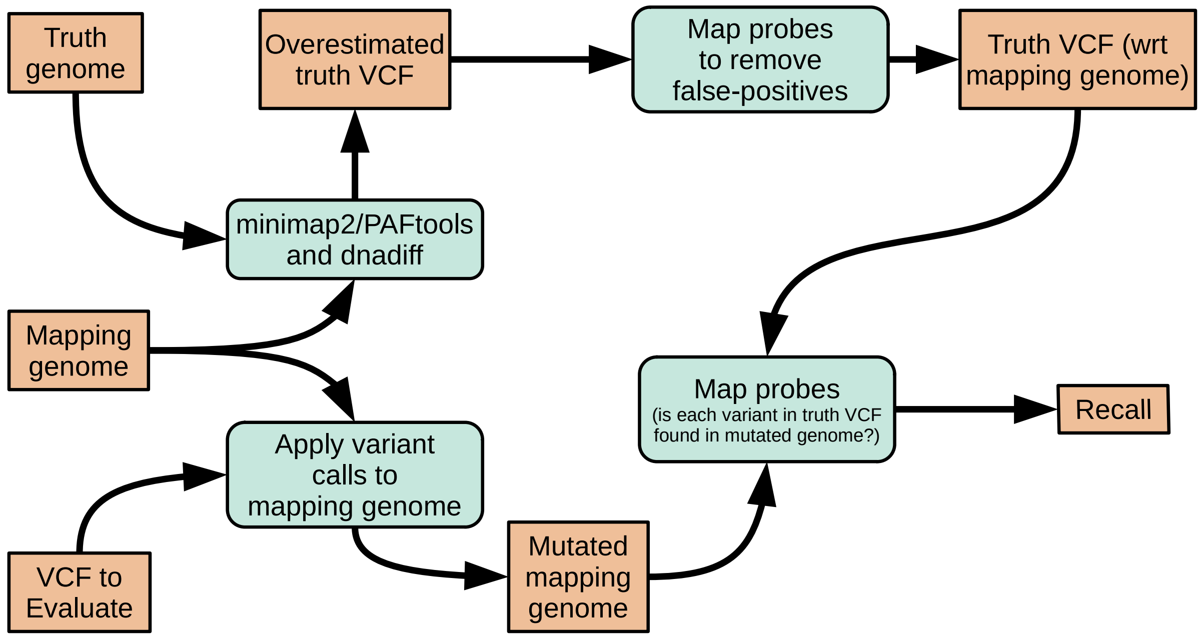


**Supplementary Figure 8:** Recall pipeline implemented by Varifier. See the Methods section in the main text for a description.

### 5 Tables

| Indel | BayesTyper | | Cortex unfiltered | | GraphTyper | | GraphTyper unfiltered | |
|---|---|---|---|---|---|---|---|---|
| Length | Precision | Recall | Precision | Recall | Precision | Recall | Precision | Recall |
| -10000 | 1.0000 | 1.0000 | 1.0000 | 1.0000 | 1.0000 | 0.9492 | 1.0000 | 0.9492 |
| -5000 | 1.0000 | 1.0000 | 1.0000 | 1.0000 | 1.0000 | 1.0000 | 1.0000 | 1.0000 |
| -2000 | 1.0000 | 1.0000 | 1.0000 | 1.0000 | 1.0000 | 1.0000 | 1.0000 | 1.0000 |
| -1000 | 1.0000 | 1.0000 | 1.0000 | 1.0000 | 1.0000 | 1.0000 | 1.0000 | 1.0000 |
| -500 | 1.0000 | 0.9946 | 1.0000 | 0.9946 | 1.0000 | 0.9946 | 1.0000 | 0.9946 |
| -100 | 1.0000 | 1.0000 | 1.0000 | 1.0000 | 1.0000 | 1.0000 | 1.0000 | 1.0000 |
| -50 | 1.0000 | 1.0000 | 1.0000 | 0.9974 | 1.0000 | 1.0000 | 1.0000 | 1.0000 |
| -20 | 1.0000 | 1.0000 | 1.0000 | 0.9973 | 1.0000 | 1.0000 | 1.0000 | 1.0000 |
| -10 | 1.0000 | 1.0000 | 1.0000 | 0.9953 | 1.0000 | 1.0000 | 1.0000 | 1.0000 |
| -5 | 1.0000 | 1.0000 | 1.0000 | 0.9956 | 1.0000 | 1.0000 | 1.0000 | 1.0000 |
| -4 | 1.0000 | 1.0000 | 1.0000 | 0.9951 | 1.0000 | 1.0000 | 1.0000 | 1.0000 |
| -3 | 1.0000 | 1.0000 | 1.0000 | 0.9953 | 1.0000 | 1.0000 | 1.0000 | 1.0000 |
| -2 | 1.0000 | 0.9998 | 1.0000 | 0.9956 | 1.0000 | 1.0000 | 1.0000 | 1.0000 |
| -1 | 1.0000 | 1.0000 | 1.0000 | 0.9949 | 1.0000 | 1.0000 | 1.0000 | 1.0000 |
| SNP | 1.0000 | 1.0000 | 1.0000 | 0.9943 | 1.0000 | 1.0000 | 1.0000 | 1.0000 |
| 1 | 1.0000 | 0.9998 | 1.0000 | 0.9946 | 1.0000 | 1.0000 | 1.0000 | 1.0000 |
| 2 | 1.0000 | 1.0000 | 1.0000 | 0.9946 | 1.0000 | 1.0000 | 1.0000 | 1.0000 |
| 3 | 1.0000 | 1.0000 | 1.0000 | 0.9944 | 1.0000 | 1.0000 | 1.0000 | 1.0000 |
| 4 | 1.0000 | 0.9995 | 1.0000 | 0.9944 | 1.0000 | 0.9998 | 1.0000 | 0.9998 |
| 5 | 1.0000 | 1.0000 | 1.0000 | 0.9941 | 1.0000 | 1.0000 | 1.0000 | 1.0000 |
| 10 | 1.0000 | 0.9998 | 1.0000 | 0.9944 | 1.0000 | 1.0000 | 1.0000 | 1.0000 |
| 20 | 1.0000 | 1.0000 | 1.0000 | 0.9946 | 1.0000 | 1.0000 | 1.0000 | 1.0000 |
| 50 | 1.0000 | 0.9944 | 1.0000 | 0.9944 | 1.0000 | 0.9944 | 1.0000 | 0.9944 |
| 100 | 1.0000 | 0.9902 | 1.0000 | 0.9902 | 1.0000 | 0.9902 | 1.0000 | 0.9902 |
| 500 | 1.0000 | 0.9900 | 1.0000 | 0.9900 | 1.0000 | 0.9851 | 1.0000 | 0.9900 |
| 1000 | 1.0000 | 0.9851 | 1.0000 | 0.9851 | 1.0000 | 0.9801 | 1.0000 | 0.9851 |
| 2000 | 1.0000 | 0.9900 | 1.0000 | 0.9900 | 1.0000 | 0.9900 | 1.0000 | 0.9900 |
| 5000 | 1.0000 | 0.9801 | 1.0000 | 0.9801 | 1.0000 | 0.9801 | 1.0000 | 0.9801 |
| 10000 | 1.0000 | 0.9652 | 1.0000 | 0.9652 | 1.0000 | 0.9652 | 1.0000 | 0.9652 |

| Indel | GraphTyper sv | | GraphTyper sv unfiltered | | Minos | | Minos unfiltered | | SAMtools unfiltered | |
|---|---|---|---|---|---|---|---|---|---|---|
| Length | Precision | Recall | Precision | Recall | Precision | Recall | Precision | Recall | Precision | Recall |
| -10000 | 1.0000 | 1.0000 | 1.0000 | 1.0000 | 1.0000 | 1.0000 | 1.0000 | 1.0000 | 1.0000 | 0.8983 |
| -5000 | 1.0000 | 1.0000 | 0.9906 | 1.0000 | 1.0000 | 1.0000 | 1.0000 | 1.0000 | 1.0000 | 0.8544 |
| -2000 | 1.0000 | 1.0000 | 1.0000 | 1.0000 | 1.0000 | 1.0000 | 1.0000 | 1.0000 | 1.0000 | 0.8701 |
| -1000 | 1.0000 | 1.0000 | 1.0000 | 1.0000 | 1.0000 | 1.0000 | 1.0000 | 1.0000 | 1.0000 | 0.8728 |
| -500 | 1.0000 | 0.9946 | 1.0000 | 0.9946 | 1.0000 | 0.9946 | 1.0000 | 0.9946 | 1.0000 | 0.8424 |
| -100 | 1.0000 | 1.0000 | 1.0000 | 1.0000 | 1.0000 | 1.0000 | 1.0000 | 1.0000 | 1.0000 | 0.5975 |
| -50 | 1.0000 | 1.0000 | 0.9855 | 0.9992 | 1.0000 | 0.9984 | 1.0000 | 1.0000 | 0.9869 | 0.9982 |
| -20 | 1.0000 | 1.0000 | 1.0000 | 1.0000 | 1.0000 | 0.9928 | 1.0000 | 1.0000 | 0.9504 | 0.9926 |
| -10 | 1.0000 | 1.0000 | 1.0000 | 1.0000 | 1.0000 | 0.9943 | 1.0000 | 1.0000 | 0.9530 | 0.9929 |
| -5 | 1.0000 | 1.0000 | 1.0000 | 1.0000 | 1.0000 | 0.9924 | 1.0000 | 1.0000 | 0.9558 | 0.9914 |
| -4 | 1.0000 | 1.0000 | 1.0000 | 1.0000 | 1.0000 | 0.9961 | 1.0000 | 1.0000 | 0.9606 | 0.9946 |
| -3 | 1.0000 | 1.0000 | 1.0000 | 1.0000 | 1.0000 | 0.9917 | 1.0000 | 1.0000 | 0.9690 | 0.9963 |
| -2 | 1.0000 | 1.0000 | 1.0000 | 1.0000 | 1.0000 | 0.9951 | 1.0000 | 1.0000 | 0.9683 | 0.9943 |
| -1 | 1.0000 | 1.0000 | 1.0000 | 1.0000 | 1.0000 | 0.9924 | 1.0000 | 1.0000 | 0.9751 | 0.9998 |
| SNP | 1.0000 | 1.0000 | 1.0000 | 1.0000 | 1.0000 | 0.9968 | 1.0000 | 1.0000 | 1.0000 | 1.0000 |
| 1 | 1.0000 | 1.0000 | 1.0000 | 1.0000 | 1.0000 | 0.9927 | 1.0000 | 1.0000 | 0.9599 | 0.9914 |
| 2 | 1.0000 | 1.0000 | 1.0000 | 1.0000 | 1.0000 | 0.9917 | 1.0000 | 1.0000 | 0.9112 | 0.9738 |
| 3 | 1.0000 | 1.0000 | 1.0000 | 1.0000 | 1.0000 | 0.9931 | 1.0000 | 1.0000 | 0.9101 | 0.9684 |
| 4 | 1.0000 | 0.9998 | 1.0000 | 0.9998 | 1.0000 | 0.9927 | 1.0000 | 0.9998 | 0.8740 | 0.9711 |
| 5 | 1.0000 | 1.0000 | 1.0000 | 1.0000 | 1.0000 | 0.9934 | 1.0000 | 1.0000 | 0.8873 | 0.9775 |
| 10 | 1.0000 | 1.0000 | 1.0000 | 1.0000 | 1.0000 | 0.9961 | 1.0000 | 1.0000 | 0.9338 | 0.9892 |
| 20 | 1.0000 | 1.0000 | 1.0000 | 1.0000 | 1.0000 | 0.9973 | 1.0000 | 1.0000 | 0.9766 | 0.9983 |
| 50 | 1.0000 | 0.9944 | 1.0000 | 0.9944 | 1.0000 | 0.9944 | 1.0000 | 0.9944 | 0.8689 | 0.0005 |
| 100 | 1.0000 | 0.9902 | 1.0000 | 0.9902 | 1.0000 | 0.9902 | 1.0000 | 0.9902 | 1.0000 | 0.0009 |
| 500 | 1.0000 | 0.9900 | 1.0000 | 0.9900 | 1.0000 | 0.9900 | 1.0000 | 0.9900 | 0.0000 | 0.0000 |
| 1000 | 1.0000 | 0.9851 | 1.0000 | 0.9851 | 1.0000 | 0.9801 | 1.0000 | 0.9851 | 0.0000 | 0.0000 |
| 2000 | 1.0000 | 0.9900 | 1.0000 | 0.9900 | 1.0000 | 0.9801 | 1.0000 | 0.9900 | 0.0000 | 0.0000 |
| 5000 | 1.0000 | 0.9801 | 1.0000 | 0.9801 | 1.0000 | 0.9751 | 1.0000 | 0.9801 | 0.0000 | 0.0000 |
| 10000 | 1.0000 | 0.9652 | 1.0000 | 0.9652 | 1.0000 | 0.9652 | 1.0000 | 0.9652 | 0.0000 | 0.0000 |

**Supplementary Table 1:** Results of calling simulated SNPs, insertions and deletions in the *M. tuberculosis* genome.

| Total length | SNPs | Insertions | Deletions | Indel length | Tool | Precision | Recall | F-score |
|---|---|---|---|---|---|---|---|---|
| 10 | 3 | 0 | 0 | 0 | BayesTyper | 0.9999 | 0.9999 | 0.9999 |
| | | | | | GraphTyper | 1.0000 | 0.9998 | 0.9999 |
| | | | | | GraphTyper unfiltered | 1.0000 | 0.9998 | 0.9999 |
| | | | | | GraphTyper sv | 1.0000 | 0.9998 | 0.9999 |
| | | | | | GraphTyper sv unfiltered | 1.0000 | 0.9998 | 0.9999 |
| | | | | | Minos | 1.0000 | 0.9979 | 0.9989 |
| | | | | | Minos unfiltered | 1.0000 | 1.0000 | 1.0000 |
| 10 | 3 | 1 | 1 | 2 | BayesTyper | 0.9981 | 0.9967 | 0.9974 |
| | | | | | GraphTyper | 0.9983 | 0.9969 | 0.9976 |
| | | | | | GraphTyper unfiltered | 0.9983 | 0.9975 | 0.9979 |
| | | | | | GraphTyper sv | 1.0000 | 0.9981 | 0.9991 |
| | | | | | GraphTyper sv unfiltered | 1.0000 | 0.9985 | 0.9993 |
| | | | | | Minos | 1.0000 | 0.9871 | 0.9935 |
| | | | | | Minos unfiltered | 0.9989 | 0.9935 | 0.9962 |
| 50 | 10 | 1 | 1 | 3 | BayesTyper | 1.0000 | 0.9953 | 0.9977 |
| | | | | | GraphTyper | 0.9998 | 0.9958 | 0.9978 |
| | | | | | GraphTyper unfiltered | 0.9997 | 0.9972 | 0.9984 |
| | | | | | GraphTyper sv | 1.0000 | 0.3689 | 0.5390 |
| | | | | | GraphTyper sv unfiltered | 0.9998 | 0.3691 | 0.5392 |
| | | | | | Minos | 1.0000 | 0.9911 | 0.9955 |
| | | | | | Minos unfiltered | 0.9996 | 0.9961 | 0.9979 |
| 50 | 20 | 3 | 3 | 4 | BayesTyper | 1.0000 | 0.9947 | 0.9973 |
| | | | | | GraphTyper | 1.0000 | 0.9947 | 0.9973 |
| | | | | | GraphTyper unfiltered | 1.0000 | 0.9947 | 0.9973 |
| | | | | | GraphTyper sv | 1.0000 | 0.9918 | 0.9959 |
| | | | | | GraphTyper sv unfiltered | 1.0000 | 0.9918 | 0.9959 |
| | | | | | Minos | 1.0000 | 0.9877 | 0.9938 |
| | | | | | Minos unfiltered | 1.0000 | 0.9945 | 0.9972 |

**Supplementary Table 2:** Results of calling simulated complex variants in the *M. tuberculosis* genome. Total length is the length of the ref allele. Columns 2–5 show the number of each type of variant added to make the complex variant. For example, the first set of variants was 3 SNPs in a window of 10bp, and no insertions or deletions. The second set was 3 SNPs, one insertion of 2bp, and one deletion of 2bp in a window of 10bp.

| Species | Number of samples | Reference genome | Tool | Mean precision | Mean recall | Mean F-score |
|---|---|---|---|---|---|---|
| *M. tuberculosis* | 17 | H37Rv | BayesTyper | 0.9995 | 0.9217 | 0.9579 |
| | | | GraphTyper | 0.9997 | 0.8938 | 0.9422 |
| | | | GraphTyper unfiltered | 0.9996 | 0.9254 | 0.9599 |
| | | | GraphTyper sv | 0.9407 | 0.8490 | 0.8889 |
| | | | GraphTyper sv unfiltered | 0.9993 | 0.8579 | 0.9000 |
| | | | Minos | 0.9997 | 0.9181 | 0.9559 |
| | | | Minos unfiltered | 0.9965 | 0.9230 | 0.9571 |
| *S. aureus* | 28 | TW20 | BayesTyper | 0.9984 | 0.8669 | 0.9279 |
| | | | GraphTyper | 0.9990 | 0.7530 | 0.8545 |
| | | | GraphTyper unfiltered | 0.9981 | 0.8785 | 0.9343 |
| | | | GraphTyper sv | 0.9963 | 0.5556 | 0.7128 |
| | | | GraphTyper sv unfiltered | 0.9956 | 0.5633 | 0.7189 |
| | | | Minos | 0.9988 | 0.8786 | 0.9347 |
| | | | Minos unfiltered | 0.9946 | 0.8953 | 0.9422 |
| | | USA300 | BayesTyper | 0.9993 | 0.8671 | 0.9283 |
| | | | GraphTyper | 0.9995 | 0.7506 | 0.8534 |
| | | | GraphTyper unfiltered | 0.9988 | 0.8829 | 0.9371 |
| | | | GraphTyper sv | 0.9982 | 0.5753 | 0.7284 |
| | | | GraphTyper sv unfiltered | 0.9972 | 0.5832 | 0.7345 |
| | | | Minos | 0.9994 | 0.8792 | 0.9353 |
| | | | Minos unfiltered | 0.9948 | 0.8970 | 0.9432 |
| *K. pneumoniae* | 17 | GCF_000784945.1 | BayesTyper | 0.9990 | 0.9052 | 0.9495 |
| | | | GraphTyper | 0.9999 | 0.9063 | 0.9505 |
| | | | GraphTyper unfiltered | 0.9997 | 0.9198 | 0.9578 |
| | | | GraphTyper sv | 0.9991 | 0.7575 | 0.8614 |
| | | | GraphTyper sv unfiltered | 0.9988 | 0.7663 | 0.8671 |
| | | | Minos | 0.9999 | 0.9143 | 0.9550 |
| | | | Minos unfiltered | 0.9974 | 0.9252 | 0.9597 |
| | | GCF_001952915.1 | BayesTyper | 0.9995 | 0.8800 | 0.9346 |
| | | | GraphTyper | 0.9999 | 0.8788 | 0.9340 |
| | | | GraphTyper unfiltered | 0.9994 | 0.8899 | 0.9402 |
| | | | GraphTyper sv | 0.9995 | 0.7092 | 0.8280 |
| | | | GraphTyper sv unfiltered | 0.9991 | 0.7181 | 0.8341 |
| | | | Minos | 0.9994 | 0.8922 | 0.9417 |
| | | | Minos unfiltered | 0.9972 | 0.8976 | 0.9437 |
| | | GCF_003073315.1 | BayesTyper | 0.9996 | 0.9267 | 0.9617 |
| | | | GraphTyper | 0.9999 | 0.9297 | 0.9634 |
| | | | GraphTyper unfiltered | 0.9997 | 0.9395 | 0.9686 |
| | | | GraphTyper sv | 0.9990 | 0.4820 | 0.6502 |
| | | | GraphTyper sv unfiltered | 0.9986 | 0.4897 | 0.6570 |
| | | | Minos | 0.9998 | 0.9367 | 0.9672 |
| | | | Minos unfiltered | 0.9987 | 0.9445 | 0.9708 |
| | | GCF_003076555.1 | BayesTyper | 0.9994 | 0.9397 | 0.9686 |
| | | | GraphTyper | 0.9999 | 0.9387 | 0.9683 |
| | | | GraphTyper unfiltered | 0.9997 | 0.9465 | 0.9723 |
| | | | GraphTyper sv | 0.9996 | 0.7968 | 0.8866 |
| | | | GraphTyper sv unfiltered | 0.9993 | 0.8010 | 0.8891 |
| | | | Minos | 0.9999 | 0.9438 | 0.9710 |
| | | | Minos unfiltered | 0.9975 | 0.9533 | 0.9748 |
| | | GCF_011006575.1 | BayesTyper | 0.9995 | 0.9078 | 0.9511 |
| | | | GraphTyper | 0.9999 | 0.9075 | 0.9513 |
| | | | GraphTyper unfiltered | 0.9997 | 0.9208 | 0.9585 |
| | | | GraphTyper sv | 0.9994 | 0.7679 | 0.8684 |
| | | | GraphTyper sv unfiltered | 0.9993 | 0.7726 | 0.8713 |
| | | | Minos | 0.9998 | 0.9238 | 0.9602 |
| | | | Minos unfiltered | 0.9974 | 0.9296 | 0.9622 |

**Supplementary Table 3:** Results of all tools evaluated on the empirical bacteria data set. Results are using the default filter of each tool, which means only taking VCF records where the `FILTER` column was equal to `PASS`. The "unfiltered" results are from ignoring the `FILTER` column and using all records.

| Dataset | Tool | Median run time (m) | Median peak RAM (GB) |
|---|---|---|---|
| Simulated | BayesTyper | 4.30 | 3.29 |
| | GraphTyper | 1.58 | 0.03 |
| | Minos | 2.62 | 0.42 |
| *M. tuberculosis* | BayesTyper | 2.91 | 3.29 |
| | GraphTyper | 1.40 | 0.03 |
| | Minos | 2.57 | 0.40 |
| *S. aureus* | BayesTyper | 10.58 | 3.33 |
| | GraphTyper | 1.67 | 0.07 |
| | Minos | 5.17 | 0.76 |
| *K. pneumoniae* | BayesTyper | 12.09 | 4.06 |
| | GraphTyper | 2.37 | 0.13 |
| | Minos | 6.01 | 2.44 |

**Supplementary Table 4:** Run time and memory summary for each tool on each data set. Values are taken from the output of the Unix command `time -v`.

| Tool | Per-sample | | Joint genotype | | | Change | |
|---|---|---|---|---|---|---|---|
| | Precision | Recall | Precision (non-ref) | Precision (all) | Recall | Precision | Recall |
| BayesTyper | 0.9995 | 0.9217 | 0.9982 | 0.9287 | 0.9225 | -0.0013 | 0.0007 |
| GraphTyper | 0.9997 | 0.8938 | 0.9998 | 0.8885 | 0.8519 | 0.0001 | -0.0419 |
| Minos | 0.9997 | 0.9181 | 0.9998 | 0.9995 | 0.9126 | 0.0001 | -0.0055 |

**Supplementary Table 5:** Change in accuracy before and after running the joint genotyping pipeline on the Walker 2013 *M. tuberculosis* data. Joint genotyped precision is calculated in two ways: using just the non-reference allele calls, and using all calls. Note that this does not apply to the recall because in that case we only look for non-reference calls, and so including reference calls has no effect.

| Data set | Task | Run time (m) | Max RAM (MB) |
|---|---|---|---|
| Mykrobe | VCF merge | 1.31 | 1744 |
| | VCF cluster | 0.63 | 7340 |
| | Gramtools build | 8.15 | 5517 |
| | Minos[1] | 85.9 | 1756 |
| | VCF merge chunks[2] | 22.97 | 505 |
| | VCF final merge | 130 | 63 |
| CRyPTIC | VCF merge | 1.56 | 2825 |
| | VCF cluster | 1.06 | 7470 |
| | Gramtools build | 13.32 | 5505 |
| | Minos[1] | 86.36 | 1769 |
| | VCF merge chunks[2] | 20.7 | 108 |
| | VCF final merge | 42.33 | 70 |

**Supplementary Table 6:** Joint genotyping Minos memory usage and run times. Values were taken from the LSF reports for each Nextflow task ("Max Memory" and "Run time"). The first three stages VCF merge, VCF cluster, and Gramtools build used 10, 20 and 20 CPUs respectively. The run times are quoted here, not total CPU time.
[1]Minos is run on each sample in parallel. The mean run time per sample and maximum RAM across all samples is shown.
[2]This process in an intermediate stage used when producing the final merged VCF file. It merges batches of 300 VCF files each into one VCF file. The merged VCF files are then input to the last task VCF final merge. The mean run time for each batch and maximum RAM across all batches is shown.

| Position | Ref | Alt | CRyPTIC all | | CRyPTIC high qual | | Mykrobe | |
|---|---|---|---|---|---|---|---|---|
| | | | Res | Sus | Res | Sus | Res | Sus |
| 426 | GTS | G | 2 | 0 | 1 | 0 | 0 | 0 |
| 427 | T | A | 0 | 1 | 0 | 1 | 0 | 0 |
| 427 | T | G | 0 | 1 | 0 | 1 | 0 | 0 |
| 427 | T | I | 1 | 1 | 0 | 1 | 2 | 0 |
| 427 | T | N | 0 | 1 | 0 | 1 | 0 | 0 |
| 428 | S | G | 1 | 1 | 1 | 0 | 1 | 0 |
| 428 | S | I | 1 | 0 | 0 | 0 | 0 | 0 |
| 428 | S | R | 1 | 0 | 1 | 0 | 3 | 0 |
| 428 | S | T | 0 | 0 | 0 | 0 | 1 | 0 |
| 429 | Q | H | 2 | 0 | 1 | 0 | 1 | 0 |
| 429 | Q | L | 7 | 0 | 7 | 0 | 0 | 0 |
| 429 | Q | P | 1 | 0 | 1 | 0 | 0 | 0 |
| 430 | L | P | 38 | 67 | 22 | 45 | 10 | 9* |
| 430 | L | R | 7 | 2 | 5 | 1 | 4 | 0 |
| 430 | LSQFM | L | 1 | 0 | 1 | 0 | 0 | 0 |
| 431 | S | G | 0 | 0 | 0 | 0 | 1 | 0 |
| 431 | S | R | 1 | 0 | 1 | 0 | 0 | 0 |
| 431 | S | SR | 1 | 0 | 1 | 0 | 1 | 0 |
| 431 | SQ | S | 0 | 0 | 0 | 0 | 1 | 0 |
| 431 | SQF | S | 0 | 0 | 0 | 0 | 1 | 0 |
| 431 | SQFM | S | 0 | 0 | 0 | 0 | 2 | 0 |
| 431 | SQFMD | S | 2 | 0 | 2 | 0 | 0 | 0 |
| 432 | Q | E | 1 | 0 | 1 | 0 | 0 | 0 |
| 432 | Q | H | 4 | 0 | 3 | 0 | 1 | 0 |
| 432 | Q | K | 12 | 0 | 9 | 0 | 4 | 0 |
| 432 | Q | L | 8 | 1 | 6 | 1 | 1 | 0 |
| 432 | Q | P | 12 | 1 | 11 | 0 | 5 | 0 |
| 433 | F | FF | 9 | 0 | 8 | 0 | 0 | 0 |
| 433 | FMD | F | 1 | 0 | 1 | 0 | 1 | 0 |
| 434 | M | I | 10 | 1 | 4 | 0 | 4 | 1 |
| 434 | M | MV | 0 | 0 | 0 | 0 | 1 | 0 |
| 434 | M | V | 1 | 0 | 0 | 0 | 0 | 0 |
| 434 | MD | M | 0 | 1 | 0 | 1 | 1 | 0 |
| 434 | MDQNNP | M | 1 | 0 | 1 | 0 | 0 | 0 |
| 435 | D | A | 3 | 2 | 1 | 0 | 1 | 1 |
| 435 | D | E | 2 | 0 | 1 | 0 | 0 | 0 |
| 435 | D | F | 21 | 2 | 13 | 1 | 5 | 0 |
| 435 | D | G | 50 | 6 | 33 | 3 | 5 | 0 |
| 435 | D | H | 1 | 0 | 1 | 0 | 2 | 0 |
| 435 | D | L | 2 | 0 | 0 | 0 | 0 | 0 |
| 435 | D | V | 418 | 20 | 335 | 10 | 79 | 0 |
| 435 | D | Y | 97 | 21 | 44 | 14 | 24 | 5* |
| 435 | DQ | D | 1 | 0 | 1 | 0 | 0 | 0 |
| 436 | Q | P | 0 | 0 | 0 | 0 | 1 | 0 |
| 436 | QN | Q | 2 | 2 | 0 | 2 | 2 | 0 |
| 437 | N | D | 7 | 1 | 3 | 1 | 2 | 0 |
| 437 | N | H | 3 | 0 | 3 | 0 | 0 | 0 |
| 437 | N | I | 1 | 0 | 1 | 0 | 0 | 0 |
| 437 | N | S | 0 | 2 | 0 | 2 | 1 | 0 |
| 439 | P | A | 1 | 0 | 1 | 0 | 0 | 0 |
| 439 | P | L | 0 | 1 | 0 | 1 | 0 | 0 |
| 441 | S | A | 1 | 0 | 1 | 0 | 0 | 0 |
| 441 | S | L | 7 | 1 | 6 | 0 | 4 | 0 |
| 441 | S | M | 0 | 0 | 0 | 0 | 1 | 0 |
| 441 | S | Q | 2 | 0 | 1 | 0 | 0 | 0 |
| 441 | S | V | 1 | 0 | 0 | 0 | 0 | 0 |
| 441 | S | W | 1 | 0 | 0 | 0 | 0 | 0 |
| 442 | G | E | 1 | 0 | 1 | 0 | 0 | 0 |
| 444 | T | S | 0 | 1 | 0 | 0 | 0 | 0 |
| 444 | T | TT | 0 | 0 | 0 | 0 | 1 | 0 |
| 444 | THK | T | 0 | 0 | 0 | 0 | 1 | 0 |
| 445 | H | C | 15 | 1 | 6 | 0 | 5 | 0 |
| 445 | H | D | 184 | 6 | 86 | 1 | 35 | 0 |
| 445 | H | F | 1 | 0 | 1 | 0 | 0 | 0 |
| 445 | H | G | 5 | 1 | 3 | 1 | 0 | 0 |
| 445 | H | L | 54 | 10 | 21 | 4 | 12 | 0* |
| 445 | H | N | 11 | 32 | 6 | 16 | 4 | 2* |
| 445 | H | P | 2 | 0 | 2 | 0 | 1 | 0 |
| 445 | H | Q | 8 | 1 | 4 | 1 | 4 | 0 |
| 445 | H | R | 26 | 1 | 23 | 1 | 9 | 1 |
| 445 | H | S | 2 | 4 | 1 | 4 | 2 | 0 |
| 445 | H | Y | 122 | 3 | 104 | 2 | 74 | 1 |
| 446 | K | E | 1 | 1 | 1 | 0 | 0 | 0 |
| 446 | K | Q | 1 | 0 | 1 | 0 | 0 | 0 |
| 446 | K | R | 2 | 0 | 2 | 0 | 0 | 0 |
| 448 | R | Q | 5 | 0 | 3 | 0 | 0 | 0 |
| 449 | L | M | 7 | 0 | 7 | 0 | 0 | 0 |
| 450 | S | A | 1 | 0 | 1 | 0 | 0 | 0 |
| 450 | S | F | 35 | 3 | 31 | 2 | 6 | 0 |
| 450 | S | G | 2 | 0 | 0 | 0 | 0 | 0 |
| 450 | S | L | 2383 | 83 | 1877 | 42 | 668 | 2 |
| 450 | S | M | 2 | 0 | 2 | 0 | 0 | 0 |
| 450 | S | Q | 1 | 0 | 1 | 0 | 2 | 0 |
| 450 | S | V | 0 | 2 | 0 | 1 | 0 | 0 |
| 450 | S | W | 47 | 2 | 38 | 2 | 27 | 0 |
| 451 | A | G | 0 | 0 | 0 | 0 | 1 | 0 |
| 451 | A | V | 4 | 0 | 3 | 0 | 1 | 0 |
| 452 | L | M | 0 | 0 | 0 | 0 | 0 | 1 |
| 452 | L | P | 69 | 40 | 29 | 21 | 25 | 6* |
| 452 | L | Q | 0 | 1 | 0 | 1 | 0 | 0 |
| 452 | L | V | 1 | 0 | 1 | 0 | 0 | 0 |

**Supplementary Table 7:** Counts of resistant and susceptible phenotypes for each amino acid variant seen in the RRDR region of the *rpoB* gene for the CRyPTIC and Mykrobe data sets. Counts are shown for all CRyPTIC samples with a rifampicin phenotype (CRyPTIC all) and only for those with a high quality phenotype (CRyPTIC high qual). Asterisk (*) shows borderline resistant mutations identified by the WHO.

| Name | Illumina sample | Illumina run | Pacbio sample | Pacbio run |
|---|---|---|---|---|
| N0004 | SAMEA4791099 | ERR2704675,ERR2704696,ERR2704697 | SAMEA3358985 | ERR964408,ERR964415 |
| N0031 | SAMEA4791100 | ERR2704676 | SAMEA3358983 | ERR956960,ERR964406 |
| N0052 | SAMEA4791101 | ERR2704677,ERR2704698,ERR2704699 | SAMEA3358982 | ERR956959,ERR964405 |
| N0054 | SAMEA4791102 | ERR2704678 | SAMEA3358987 | ERR964410,ERR964417 |
| N0072 | SAMEA4791104 | ERR2704680 | SAMEA3358979 | ERR956956,ERR964402 |
| N0091 | SAMEA4791105 | ERR2704681 | SAMEA3358993 | ERR964422,ERR968288 |
| N0136 | SAMEA4791106 | ERR2704682,ERR2704700 | SAMEA3358989 | ERR964412,ERR964419 |
| N0145 | SAMEA4791107 | ERR2704683,ERR2704701,ERR2704702 | SAMEA3358981 | ERR956958,ERR964404 |
| N0153 | SAMEA5383013 | ERR3183990 | SAMEA3358980 | ERR956957,ERR964403 |
| N0155 | SAMEA4791108 | ERR2704684,ERR2704703 | SAMEA3358984 | ERR964407,ERR964414 |
| N0157 | SAMEA4791109 | ERR2704685,ERR2704704 | SAMEA3358978 | ERR956955,ERR964401 |
| N1176 | SAMEA4791110 | ERR2704686 | SAMEA3443999 | ERR1025289,ERR1036243 |
| N1177 | SAMEA5383014 | ERR3183991 | SAMEA3444000 | ERR987691,ERR999930 |
| N1202 | SAMEA4791112 | ERR2704688 | SAMEA3358994 | ERR968284,ERR968289 |
| N1216 | SAMEA4791113 | ERR2704689,ERR2704705 | SAMEA3358988 | ERR964411,ERR964418 |
| N1272 | SAMEA4791115,SAMEA4791116 | ERR2704691,ERR2704692,ERR2704707,ERR2704708 | SAMEA3358992 | ERR964421,ERR968287 |
| N1283 | SAMEA4791118 | ERR2704694,ERR2704709 | SAMEA3358990 | ERR964413,ERR968285 |

**Supplementary Table 8:** *M. tuberculosis* accessions.

| Name | Illumina sample | Illumina run | Assembly |
|---|---|---|---|
| KSB1_9D | SAMN07211282 | SRR5665580 | GCF_002753165.1 |
| INF157 | SAMEA3357077 | ERR1023800 | GCF_002753075.1 |
| INF158 | SAMEA3357080 | ERR1023801 | GCF_002753055.1 |
| INF274 | SAMEA3357268 | ERR1008735 | GCF_002753605.1 |
| INF278 | SAMEA3357272 | ERR1008737 | GCF_002752975.1 |
| INF042 | SAMEA3357010 | ERR1023775 | GCF_002752995.1 |
| INF059 | SAMEA3357043 | ERR1023792 | GCF_002753355.1 |
| KSB1_7J | SAMEA3357374 | ERR1023671 | GCF_002753375.1 |
| INF322 | SAMEA3357193 | ERR1008775 | GCF_002752815.1 |
| INF249 | SAMEA3357223 | ERR1008714 | GCF_002752775.1 |
| KSB1_4E | SAMEA3357325 | ERR1023628 | GCF_002754835.1 |
| KSB1_7E | SAMEA3357328 | ERR1023631 | GCF_002752905.1 |
| KSB1_10J | SAMEA3357381 | ERR1023674 | GCF_002753035.1 |
| KSB2_1B | SAMEA3357405 | ERR1023685 | GCF_002753015.1 |
| INF163 | SAMN07211279 | SRR5665579 | GCF_002753405.1 |
| INF164 | SAMN07211280 | SRR5665578 | GCF_002753555.1 |
| KSB1_5D | SAMN06112197 | SRR5082456 | GCF_002741685.1 |

**Supplementary Table 9:** *K. pneumoniae* accessions.

| Name | Illumina sample | Illumina run |
|---|---|---|
| C00001076 | SAMEA2335070 | ERR410035 |
| C00001108 | SAMEA2335102 | ERR410067 |
| C00001130 | SAMEA2335124 | ERR410089 |
| C00001155 | SAMEA2335149 | ERR410114 |
| C00001205 | SAMEA2340393 | ERR418876 |
| C00001284 | SAMEA2335171 | ERR410136 |
| C00012767 | SAMEA2340499 | ERR418982 |
| C00012779 | SAMEA2340511 | ERR418994 |
| C00012783 | SAMEA2340515 | ERR418998 |
| C00012789 | SAMEA2340521 | ERR419004 |
| C00012812 | SAMEA2340544 | ERR419027 |
| C00013168 | SAMEA2340580 | ERR419063 |
| C00013231 | SAMEA2340643 | ERR419126 |
| C00013240 | SAMEA2340651 | ERR419134 |
| C00013352 | SAMEA2340702 | ERR419185 |
| C00013361 | SAMEA2340711 | ERR419194 |
| C00013369 | SAMEA2340719 | ERR419202 |
| C00013375 | SAMEA2340725 | ERR419208 |
| C00013377 | SAMEA2340727 | ERR419210 |
| C00013388 | SAMEA2340738 | ERR419221 |
| C00013390 | SAMEA2340740 | ERR419223 |
| C00013391 | SAMEA2340741 | ERR419224 |
| C00013399 | SAMEA2340749 | ERR419232 |
| C00013402 | SAMEA2340752 | ERR419235 |
| C00013405 | SAMEA2340755 | ERR419238 |
| C00013406 | SAMEA2340756 | ERR419239 |
| C00013407 | SAMEA2340757 | ERR419240 |
| C00001125 | SAMEA2335119 | ERR410084 |

**Supplementary Table 10:** *S. aureus* accessions.
